# Input-specific localization of NMDA receptor GluN2 subunits in thalamocortical neurons

**DOI:** 10.1101/2024.08.23.607324

**Authors:** Mackenzie A. Topolski, Brian L. Gilmore, Rabeya Khondaker, Juliana A. Michniak, Carleigh Studtmann, Yang Chen, Gwen N. Wagner, Aaron E. Pozo-Aranda, Shannon Farris, Sharon A. Swanger

**Affiliations:** Fralin Biomedical Research Institute at VTC, Virginia Tech, Roanoke, VA, USA; Graduate Program in Translational Biology, Medicine, and Health, Virginia Tech, Blacksburg, VA, USA; Department of Biological Systems Engineering, Virginia Tech, Blacksburg, VA, USA; School of Neuroscience, Virginia Tech, Blacksburg, VA, USA; Department of Biomedical Sciences and Pathobiology, Virginia-Maryland College of Veterinary Medicine, Virginia Tech, Blacksburg, VA, USA; Department of Internal Medicine, Virginia Tech Carilion School of Medicine, Roanoke, VA, USA

**Keywords:** NMDA receptor, GluN2, glutamatergic synapse, postsynaptic receptor, synapse diversity, thalamocortical neuron, thalamus

## Abstract

Molecular and functional diversity among synapses is generated, in part, by differential expression of neurotransmitter receptors and their associated protein complexes. *N*-methyl-*D*-aspartate receptors (NMDARs) are tetrameric ionotropic glutamate receptors that most often comprise two GluN1 and two GluN2 subunits. NMDARs generate functionally diverse synapses across neuron populations through cell-type-specific expression patterns of GluN2 subunits (GluN2A – 2D), which have vastly different functional properties and distinct downstream signaling. Diverse NMDAR function has also been observed at anatomically distinct inputs to a single neuron population. However, the mechanisms that generate input-specific NMDAR function remain unknown as few studies have investigated subcellular GluN2 subunit localization in native brain tissue. We investigated NMDAR synaptic localization in thalamocortical (TC) neurons expressing all four GluN2 subunits. Utilizing super resolution imaging and knockout-validated antibodies, we revealed subtype- and input-specific GluN2 localization at corticothalamic (CT) versus sensory inputs to TC neurons in 4-week-old male and female C57Bl/6J mice. GluN2B was the most abundant postsynaptic subunit across all glutamatergic synapses followed by GluN2A and GluN2C, and GluN2D was localized to the fewest synapses. GluN2B was preferentially localized to CT synapses over sensory synapses, while GluN2A and GluN2C were more abundant at sensory inputs compared to CT inputs. Furthermore, postsynaptic scaffolding proteins PSD95 and SAP102 were preferentially localized with specific GluN2 subunits, and SAP102 was more abundant at sensory synapses than PSD95. This work indicates that TC neurons exhibit subtype- and input-specific localization of diverse NMDARs and associated scaffolding proteins that likely contribute to functional differences between CT and sensory synapses.

**HIGHLIGHTS:**

- NMDAR subtypes and synaptic scaffolding proteins show preferential localization at specific inputs to thalamocortical neurons.
- GluN2B was preferentially localized to corticothalamic synapses, while GluN2A and GluN2C were more abundant at sensory inputs to thalamocortical neurons.
- Colocalization between synaptic scaffolding proteins with NMDARs was GluN2 subtype-dependent.
- NMDAR subsynaptic organization in thalamocortical neurons is input- and GluN2-subtype specific.

## INTRODUCTION

Molecular heterogeneity among synapses generates functionally diverse synapses, cells, and circuits in the central nervous system (CNS). Synapse-specific molecular signatures yield distinct electrophysiological properties, signal transduction pathways, and synapse structure (Grant & Fransen, 2020). In turn, synapse diversity leads to cell-type-specific information processing capacities and circuit-specific susceptibility to disease. Glutamatergic synapses mediate excitatory neurotransmission throughout the CNS, but there is substantial functional heterogeneity generated by glutamate receptor subtypes (Frank & Grant, 2017). *N*-methyl-*D*-aspartate receptors (NMDARs) are most often composed of two GluN1 and two GluN2 subunits, of which there are four subtypes: GluN2A, GluN2B, GluN2C, and GluN2D (Hansen et al., 2021). GluN2 subunits confer NMDARs with different biophysical properties that generate distinct synaptic time courses, calcium flux, and voltage dependence. Furthermore, GluN2 subunits associate with different macromolecular complexes that can activate distinct downstream signaling and plasticity mechanisms (Paoletti et al., 2013).

Distinctive expression of the genes encoding GluN2 subunits creates molecular and functional diversity of NMDARs across different cell-types and brain regions (Akazawa et al., 1994; Monyer et al., 1994). NMDAR function also differs among anatomically distinct inputs to or from a single neuron population. This includes input-specific postsynaptic NMDAR function in the cortex, thalamus, amygdala, and spinal cord (Deleuze & Huguenard, 2016; Ferrer et al., 2018; Jung et al., 2010; Kumar & Huguenard, 2003; Lewis et al., 2022; Miyata & Imoto, 2006) as well as target-specific presynaptic NMDAR function in the hippocampus and cortex (Brasier & Feldman, 2008; Lituma et al., 2021). Notably, GluN2 contributions can also differ among synaptic inputs of the same type onto a single neuron population as shown for primary afferent synapses on spinal cord neurons (Pitcher et al., 2023). The mechanisms underlying synapse-specific NMDAR localization and function remain poorly understood, particularly in native CNS tissue. Input- and target-specific NMDAR function could result from different expression levels of all NMDARs across synapses, synapse-specific localization of NMDAR subtypes, and/or distinct mechanisms controlling NMDAR activation (e.g. synapse-specific release properties, glutamate transporter efficiency, or AMPA receptor content). Herein, we sought to determine how input-specific localization of GluN2 subunits contributed to molecular diversity among synapses in thalamocortical (TC) neurons.

TC neurons are an ideal cell population to probe NMDAR diversity as they simultaneously express all four GluN2 subtypes at appreciable levels (Phillips et al., 2019; Standaert et al., 1994). Diverse GluN2 expression has been observed across thalamic nuclei, but input-specific GluN2 function has only been shown in the somatosensory ventrobasal (VB) thalamus, which comprises the ventral posterolateral (VPL) and posteromedial (VPM) regions. TC neurons in VPM thalamus receive two main glutamatergic inputs: ascending brainstem inputs carrying somatosensory information and descending layer 6 corticothalamic (CT) inputs. GluN2B mediates synaptic transmission at both synapses during early postnatal development in 1-week-old mice, but GluN2B function is only observed at CT inputs, not sensory inputs, after 2 weeks of age (Miyata, 2007; Miyata & Imoto, 2006). GluN2B may be lost from sensory synapses or it could be localized extrasynaptically later in postnatal development making its functional contributions more difficult to detect. Electron microscopy showed that synaptic localization of GluN2B in the thalamus decreased during postnatal development, but this study did not investigate input-specific localization (Liu et al., 2004). Furthermore, the localization of GluN2A, GluN2C, and GluN2D remain unknown in TC neurons. In this study, we investigated how GluN2 subunit organization differs between CT and sensory inputs to VPM thalamus neurons. We discovered input-specific NMDAR localization as well as GluN2 subtype-specific synaptic organization including preferential localization with specific postsynaptic scaffolding proteins. These data reveal molecular diversity that likely contributes to synapse-specific NMDAR function in VPM neurons and suggest that TC neurons harbor mechanisms for localizing NMDARs in a sub-type- and input-specific manner.

## MATERIALS AND METHODS

### Animal models

Mouse studies were performed according to protocols approved by the Institutional Animal Care and Use Committee at Virginia Polytechnic Institute and State University and in accordance with the National Institutes of Health guidelines. Experiments utilized C57Bl/6J mice of both sexes aged 2 to 8 weeks that were obtained from Jackson Laboratory (RRID: IMSR_JAX:000664). Mice were housed in a 12-hour light/dark cycle with *ad libitum* access to food and water. Brain tissue from 4-week-old *Grin2a-/-* (Kadotani et al., 1996), *Grin2c-/-* (Karavanova et al., 2007), and *Grin2d-/-* (Ikeda et al., 1995) mice was generously provided by Dr. Stephen Traynelis.

### Fluorescence in situ hybridization imaging and analysis

Four-week-old C57Bl/6J mice were euthanized by isoflurane overdose. The brains were dissected, embedded in OCT, and rapidly frozen by submersion in a 2-methylbutane bath on dry ice. Twenty-micron cryosections were mounted on slides and then processed for single-molecule fluorescence in situ hybridization (FISH) according to the manufacturer instructions in the RNAscope HiPlex kit (Advanced Cell Diagnostics). Probes used included: *Grin1 (*431611-T7)*, Grin2a* (481831-T1)*, Grin2b* (417391-T2)*, Grin2c* (445581-T3)*, Grin2d* (425951-T5), and *Slc17a6 (*317011-T6) as well as manufacturer’s negative and positive controls. Whole brain section images were acquired as tiled 40x images on an Olympus IX83 widefield fluorescence microscope with an XCite light source, Hamamatsu Orca Flash 4.0 camera, and DAPI, FITC, TRITC, and Cy5 filter sets. Z-stack images (3 µm interval) for single particle quantification were acquired using a Zeiss 700 scanning confocal with 405 nm, 488 nm, 561 nm, and 647 nm laser lines and a 40X objective. Laser power and gain settings were such that the negative control had ≤ 3 particles visible per cell. Acquisition settings were held constant for all images from the same round of hybridization. Five z-planes were processed and analyzed for mRNA particle counting. Threshold values were set based on values for negative controls having ≤ 3 particles per cell. Values were held constant within each channel for all images within one hybridization round. A binary image with individual cell ROIs was made by applying a gaussian blur filter (σ = 2) to composite images with DAPI and all mRNA channels. Particles per ROI were counted from the thresholded images utilizing the *Analyze Particles* plug-in and a self-written Fiji macro.

### Western blotting

Two- to eight-week-old mice were euthanized by isoflurane overdose and transcardially perfused with ice-cold 1x phosphate-buffered saline (PBS). Brains were dissected, the cerebellum was removed, and VB thalamus tissue punches were taken from coronal slices (500 µm) made with a Leica 1200S vibratome. Tissue was snap frozen in liquid nitrogen. VB punches and cerebellum tissue were sonicated in homogenization buffer (1 mM HEPES, 320 mM sucrose, 1X mini Complete protease inhibitor cocktail, pH 7.4). Supernatants were collected after centrifugation (15 min, 17,000*xg*, 4°C), and protein concentration was measured using Pierce Rapid Gold BCA Protein Assay. Samples were prepared in 1x Bio-Rad Laemmli sample buffer (250 mM dithiothreitol), heated at 95°C for 5 min, and loaded on duplicate Bio-Rad TGX stain-free 4-15% gradient gels. Protein concentrations were normalized for each brain region across ages. Gels were imaged with the Bio-Rad ChemiDoc MP stain-free detection protocol. Proteins were transferred to Bio-Rad LF PVDF in 1x Bio-Rad Trans-blot Turbo transfer buffer (20% ethanol) using the high molecular weight program.

The blots were imaged using the stain-free blot detection protocol, allowed to dry, and then rehydrated before blocking for 1 hour in 2% non-fat milk in Tris-buffered saline (TBS). The duplicate blots were probed with GluN2C [2% milk in TBS with 0.1% Tween 20 (TBST)] or GluN2D (2% milk in TBST with 0.02% SDS; **Table 1**). All primary antibodies were incubated overnight (∼16 hr) at 4°C and washed in TBST (0.05% Tween) four times for a total of 30 minutes. Goat anti-mouse HRP-conjugated secondary antibody was diluted (1/10,000, **Table 2**) in the respective milk solutions used for primary antibodies and incubated for 1 hour at room temperature (RT). Blots were washed again before adding SuperSignal Pico PLUS western blot substrate for 5 minutes and imaged on the ChemiDoc MP (chemiluminescence, Signal Accumulation Mode protocol). The blots were stripped for 30 min with Restore PLUS before being blocked again for 1 hour in 2% milk in TBS. The blots were re-probed with GluN2A or GluN2B antibodies (**Table 1**) diluted in 2% milk in TBST (0.05% Tween) and washed as above. Goat anti-rabbit HRP-conjugated secondary antibody (1/50,000, **Table 2**) or goat anti-mouse HRP-conjugated secondary antibody (1/10,000) in 2% milk/TBS-T (0.05%) was incubated for 1 hour, and then the blots were washed and imaged. The blots were stripped and blocked again before re-probing with GluN1 antibody (**Table 1**) and goat anti-mouse HRP-conjugated secondary antibody (1/10,000) diluted in 2% milk/TBST (0.05%) followed by imaging as above.

**Table 1.**
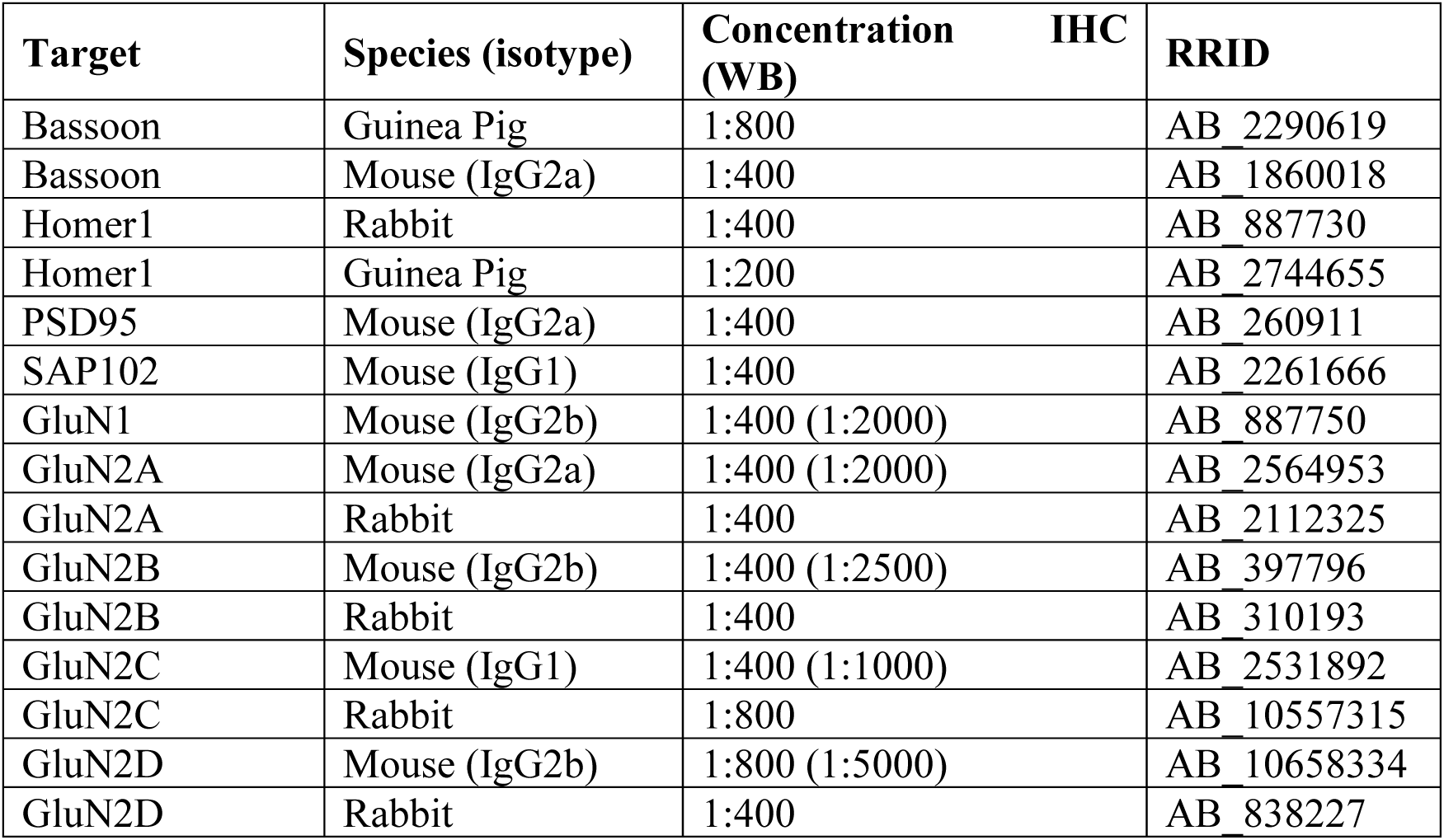
Primary antibodies.

**Table 2.**
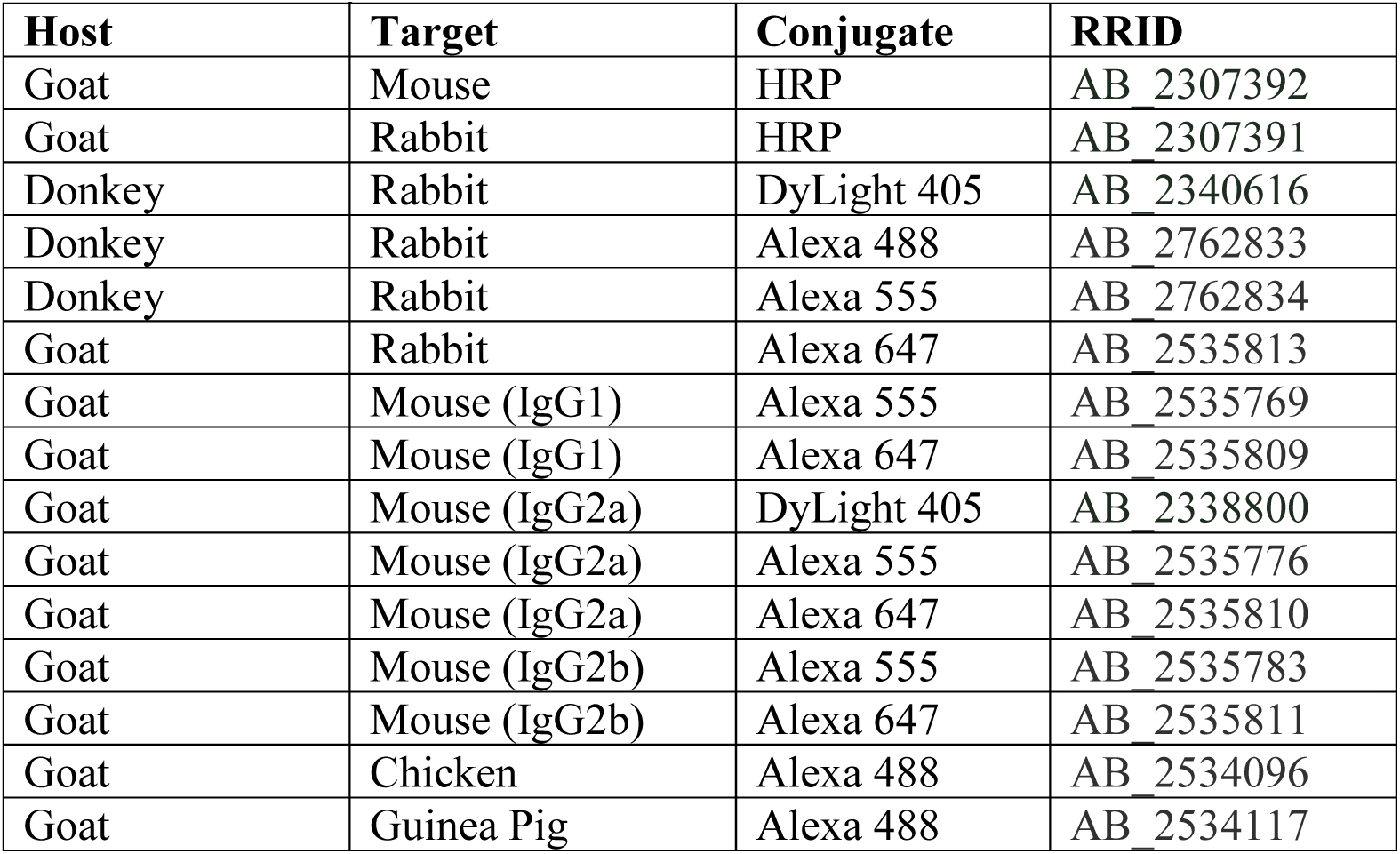
Secondary antibodies.

### Immunohistochemistry

Mice were transcardially perfused with 1x PBS followed by 4% paraformaldehyde (PFA) in 1x PBS, and the brains were post-fixed in 4% PFA for 24 hr, cryoprotected in 30% sucrose in 1x PBS, and frozen in OCT. Cryosections (20 μm) were treated with 0.8% sodium borohydride in 1x TBS, pH 7.4, at RT for 10 min, washed with 1x TBS, and then treated with 0.01 M sodium citrate buffer, pH 6.0, at 100°C for 10 min. Sections were cooled at RT for 30 min, washed with 1x TBS, and then permeabilized with 0.5% Triton X-100 in 1x TBS for 10 min. Tissue was blocked with Fab fragments (40 μg/ml) in 1x TBS for 30 min at RT, followed by blocking with 10% normal donkey serum (NDS) or normal goat serum (NGS) in 1x TBS with 0.1% Triton X-100 (TBS-TX). Tissue was incubated with primary antibodies (**Table 1**) diluted in 2% NDS or NGS in 1x TBS-TX overnight at 4°C, and then washed with 1x TBS three times for 10 min each. Secondary antibodies (**Table 2**) were diluted (1:400) with 2% NDS or NGS in 1x TBS-TX for 1 hr at RT, followed by three 10-minute washes with 1x TBS. All GluN2 antibodies were detected with Alexa 555-conjugated antibodies in each immunostaining. Bassoon was detected with Alexa 405- or 488-conjugated antibodies. Homer, PSD95, and SAP102 were detected with Alexa 488- or 647-conjugated antibodies. VGLUT1 and VGLUT2 were detected with Alexa 405-, 488-, or 647-conjugated antibodies. Coverslips were mounted with Prolong Gold mounting media, and slides were stored at RT overnight, and then at 4°C until imaging.

### Immunohistochemistry image acquisition and analysis

Whole brain sections were imaged by tiling of 10x images acquired on an Olympus IX83 microscope described above. High-resolution synapse images were acquired using a Nikon super resolution by optical reassignment (SoRa) spinning disk confocal microscope with a 60x objective and 4x SoRa disk. Fluorescence signals were excited using 405, 488, 561, and 640 nm lasers and detected using DAPI, GFP, RFP, and Cy5 filters and a Hamamatsu Back-Thinned Orca Fusion CMOS camera. Z-stacks were acquired with Nikon Elements software with a 0.15 μm interval. Three 60x images from different fields of view (FOV) within the VPM thalamus were taken for each mouse. Images were processed with Nikon Elements Blind Deconvolution algorithm. All further image processing and analysis was performed using Fiji on a single z-plane from each image series.

A nearest neighbor (NN) analysis was performed on raw images without any thresholding utilizing the MOSAIC Fiji plugin with the default values for size and kernel distance (Shivanandan et al., 2013). Interaction strength values were used to compare the degree of correlation of spatial distributions across signal pairs. The SynapseJ.ijm plugin was used to detect synapses defined as overlapping homer and bassoon, PSD95 and bassoon, or SAP102 and bassoon (Moreno Manrique et al., 2021). FindMaxima.ijm plugin was utilized to set the minimum and maximum threshold intensities and noise level for detecting each synaptic marker and NMDAR as separate particles. These values were utilized within the SynapseJ plugin to detect synapses and kept constant for each immunogen across all images from a single imaging session. To improve detection of synapses and NMDARs localized near a synapse, we dilated the signals for bassoon, homer, PSD95, and SAP102 as well as GluN1 and GluN2 subunits by 2 pixels using the median blur filter. No median blur or dilation was applied within the SynapseJ plugin. VGLUT-positive synapses were defined as homer/bassoon, PSD95/bassoon, or SAP102/bassoon pairs with bassoon signal overlapping VGLUT1 or VGLUT2 signal with at least one pixel at a maximum intensity threshold set empirically for VGLUT1 or VGLUT2 signals. Synapses with postsynaptic GluN subunits were defined as synapses with the postsynaptic marker (homer, PSD95, or SAP102) overlapping with GluN signal above a maximum intensity threshold. Synapses with presynaptic GluN were defined as synapses with bassoon signal overlapping a GluN particle, but not homer. If a GluN particle was overlapping both bassoon and a postsynaptic marker, then the GluN particle was classified as overlapping that with the shortest distance, which was the postsynaptic marker in >95% of cases. The distances between puncta were measured from the maxima. The data from three FOV were averaged for each mouse. Experimenters were blind to immunogen identity during imaging and image analysis.

### Statistical analysis

*A priori* power analyses were performed in GPower 3.1 to estimate required samples sizes given appropriate statistical tests with α = 0.05, power (1 – β) = 0.8, and a moderate effect size or effect sizes based on pilot data. Statistical analyses were performed in GraphPad (Prism). All datasets were tested for normality with the Shapiro-Wilk test and equal variances using an F test (two independent groups) or the Brown-Forsythe test (three or more groups). Most datasets were normally distributed, but we routinely saw unequal variances in our pilot data. Therefore, normal datasets were analyzed by Welch’s t-test, ANOVA with Brown-Forsythe correction, or a mixed-effects model, as appropriate. Pairwise analyses were corrected for multiple comparisons with the specific tests listed in the figure legends. Group data were plotted as mean ± *SD* in the figures, and numerical data reported in the text are mean ± *SD* or median (interquartile range, IQR), unless otherwise stated. The 95% confidence intervals (CI) of the difference between means (MD) were plotted as part of an estimation plot (two groups) or a separate plot next to the group data. The *p* values for pairwise comparisons are noted on the plots with the group data, or on the 95% CI plot when there were more pairwise comparisons made than could be clearly visualized on group data plot. Critical values and degrees of freedom (*df*) were stated in the figure legends.

## RESULTS

### Quantification of NMDAR localization at single synapses in the thalamus

To examine native NMDAR synaptic localization, we detected single glutamatergic synapses by immunolabeling bassoon (presynaptic marker) and homer (postsynaptic marker) in thalamus tissue sections (**Figure 1A**). Super resolution imaging by optical pixel reassignment (SoRa) afforded XY resolution of ∼120 nm and Z resolution of ∼300 nm (Azuma & Kei, 2015) (**Figure 1B**). We first confirmed that the spatial distributions of homer and bassoon were correlated in the thalamus utilizing a nearest neighbor (NN) analysis. If homer and bassoon are effective markers of thalamic synapses, then the NN analysis should yield a positive interaction strength value indicating their distribution is correlated. As expected, bassoon and homer interaction strength (2.35, *SD* 0.50) was significantly greater than control images with the homer signal rotated 90° (0.12, *SD* 0.46; **Figure 1C**). We performed automated synapse detection with the SynapseJ plugin that identified overlapping pairs of bassoon and homer (Moreno Manrique et al., 2021) (**Figure 1D,E**). Given the XY image resolution of 120 nm, juxtaposed presynaptic and postsynaptic puncta may not always overlap; to ensure we detected these pairs, we dilated both presynaptic and postsynaptic puncta by two pixels with a median blur filter. Utilizing this method, we detected 42% (*SD* 5) of bassoon puncta and 33% (*SD* 6) of homer puncta as glutamatergic synapses (**Figure 1F**). Less than 3% (*SD* 2) of synaptic homer puncta were overlapping more than one bassoon puncta, which supports the assertion that this method effectively identified individual synapses. Furthermore, the median distance between homer and bassoon was 193 nm (IQR 31), which is consistent with previous reports utilizing other super resolution imaging modalities (Dani et al 2010) (**Figure 1G**).

**Figure 1.**
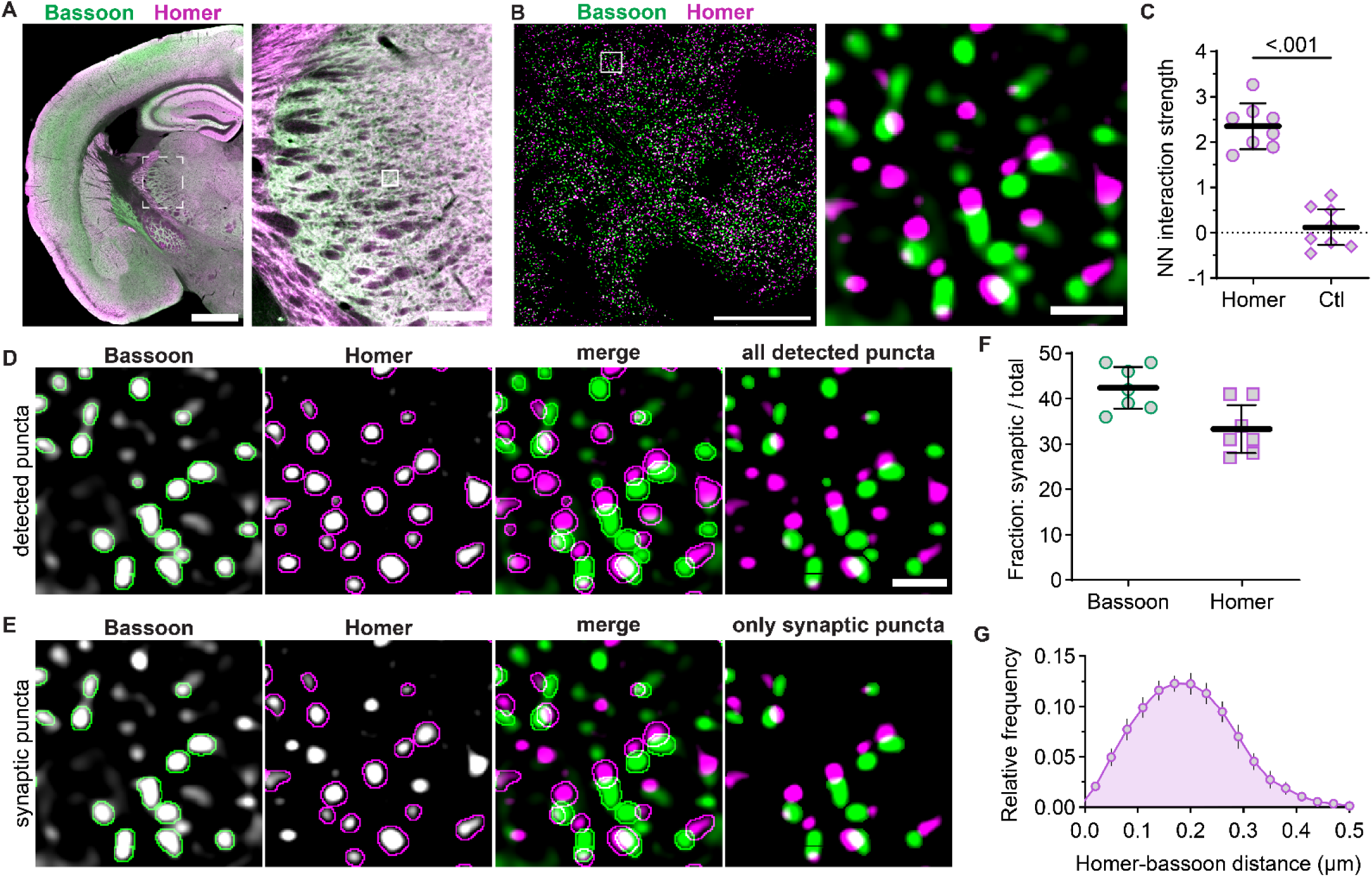
Detection of single synapses in VPM thalamus. **A.** A tiled 10x image of a mouse brain section shows homer (magenta) and bassoon (green) immunostaining with a portion of the thalamus (dashed region) shown at right. The solid box is a representative field of view (FOV) for 60x images with 4x SoRa in the VPM. **B.** A 60x image (4x SoRa) shows homer and bassoon puncta. The boxed region is shown at right to better visualize individual homer and bassoon puncta. **C**. The NN interaction strengths between homer and bassoon were determined for 60x (4x SoRa) images for each mouse using the original images (Homer) and control images with homer signals rotated 90° (Ctl). The plots show data points for individual mice with the group mean ± *SD*. Welch’s t-test, *t =* 9.173, *df* = 13.92, *n =* 8 mice; 4 male (4M) and 4 female (4F). **D.** Homer and bassoon signals from panel B are shown separately in grayscale and merged with outlines depicting all detected puncta from SynapseJ analysis. The thresholded “only detected puncta” image shows only homer and bassoon puncta analyzed for colocalization to detect synapses. **E**. Images from panel D are shown with outlines depicting only puncta identified as presynaptic bassoon and postsynaptic homer in grayscale and merged. The thresholded “only synaptic puncta” image shows only the outlined bassoon and homer puncta. **F.** The percentages of total bassoon and homer puncta identified as synaptic were plotted for each mouse with group mean ± *SD* (n = 7 mice; 3M, 4F). **G**. The distances between homer and bassoon synaptic puncta were measured and plotted as a frequency distribution for each mouse from panel F. The plot shows the group mean frequency distribution ± *SD*. Scale bars: (A) 1 mm, 200 µm; (B) 20 µm, 1 µm.

To detect native NMDARs at single synapses, we first examined GluN1, which is an obligate subunit of all NMDARs (**Figure 2A**). NMDAR function has been observed at both CT and sensory synapses, and thus, GluN1 was expected to exhibit a high degree of postsynaptic localization in VPM neurons. We performed bassoon, homer, and GluN1 co-immunostaining, acquired super resolution images, and detected homer/bassoon synapses (**Figure 2B**). We then quantified the percentage of synapses that had GluN1 overlapping homer or bassoon (**Figure 2C,D**). GluN1 overlapped with homer at 50% (*SD* 9) of synapses and bassoon at 7% (*SD* 2) of synapses (**Figure 2E,F)**. These data indicate that we can detect postsynaptic NMDARs at approximately half of VPM glutamatergic synapses and that a small proportion of VPM synapses may have presynaptic NMDARs. Postsynaptic NMDARs are likely present at more synapses than we detected as we analyzed a single z-plane with strict thresholding to limit erroneous synapse colocalization with out-of-plane objects.

**Figure 2.**
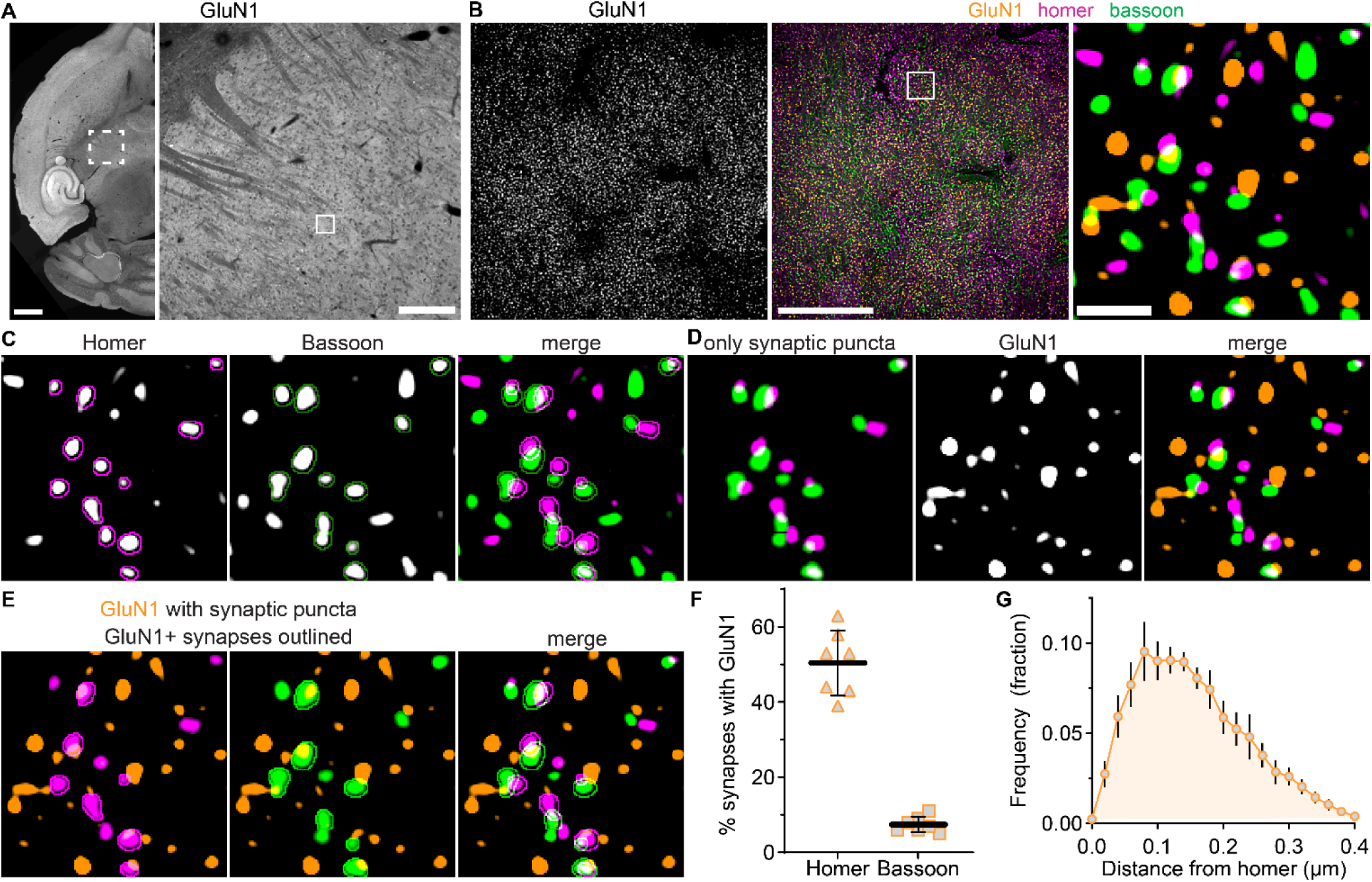
Detection and quantification of GluN1 synaptic localization. **A.** A tiled 10x image of a horizontal mouse brain section shows GluN1 immunostaining with a region of the thalamus (dashed box) shown at right. The solid box is a representative 60x (4x SoRa) FOV in the VPM. **B.** 60x (4x SoRa) images show GluN1 immunostaining and merged GluN1 (orange), homer (magenta), and bassoon (green) signals with the boxed region expanded at right. **C.** Homer and bassoon signals from panel B are shown separately in grayscale and merged with outlines depicting detected synaptic puncta. **D**. The thresholded image of homer and bassoon signals show only synaptic puncta. GluN1 signal is shown in grayscale and then merged with the “only synaptic puncta” image. **E.** GluN1 signal is shown with postsynaptic homer and presynaptic bassoon signals from panel **D** with outlines around only those synapses with GluN1 overlapping homer (GluN1+). **F**. The percentage of total synaptic puncta identified as having postsynaptic (Homer) and presynaptic (Bassoon) GluN1 were plotted (n = 7 mice; 3M, 4F) with the mean ± *SD*. **G**. The distance between each postsynaptic homer and respective GluN1 puncta was measured and plotted as a frequency distribution. The plot shows the group mean frequency distribution ± *SD.* Scale bars: (A) 1 mm, 200 µm; (B) 20 µm, 1 µm.

The median distance between postsynaptic GluN1 and homer was 161 nm (*IQR* 55) indicating that we detected NMDARs localized within the same synaptic domain as homer (**Figure 2G**). The longest distance between GluN1 and homer that was considered to be postsynaptic was 400 nm, and thus, our analysis likely includes both synaptic and extrasynaptic NMDARs. Interestingly, the frequency distribution of GluN1-homer distances was non-normal (K^2^ = 8.85; *p* = 0.012) with a broad peak and was skewed rightward. We surmised that the shape of this distribution may be due to distinct subsynaptic organization across sensory and cortical inputs and/or distinct GluN2 subsynaptic organization at one or both inputs. Analysis of GluN1 localization was used to demonstrate that we effectively detected NMDARs at single synapses. We subsequently applied this method to compare the synaptic organization of GluN2A – 2D at all glutamatergic synapses and specific synaptic inputs in the VPM thalamus.

### Synaptic localization of native NMDAR differs among GluN2 subunits

To study GluN2 synaptic localization in the VPM thalamus, we first determined the age that maximized expression of GluN2A – 2D subunits in VB thalamus tissue punches (comprising the VPL and VPM) by immunoblotting. Developmental expression of GluN2 subunits differs across brain regions and had not, to our knowledge, been quantified specifically in the thalamus. GluN2A and GluN2C expression were low at two weeks and increased through the fourth postnatal week, while GluN2B and GluN2D expression was appreciable at all ages through adulthood (**Supplemental Figure S1**). We determined that expression of all four GluN2 proteins in VB thalamus was maximized in 4-week-old mice, which were used for the remainder of this study.

We further evaluated the relative expression levels of GluN2 subunits in VPM thalamus by comparing mRNA levels within VPM neurons by single molecule FISH using HiPlex RNAscope (**Supplemental Figure S2**). *Slc17a6* (VGLUT2) mRNA was labeled to identify TC neurons. The mean mRNA particle number per cell for *Grin2a* (53 particles/cell, *SD* 3) and *Grin2b* (55, *SD* 2) were similar, but *Grin2c* mRNA levels (33, *SD* 4) were significantly lower than *Grin2a* and *Grin2b*. *Grin2d* mRNA levels (14, *SD* 1) were significantly lower than *Grin2a*, *Grin2b,* and *Grin2c*. Second, we examined GluN2 protein expression in VPM thalamus by immunolabeling GluN2A, GluN2B, GluN2C, or GluN2D in mouse brain sections (**Figure 3A**). The relative number of GluN2 protein puncta largely corresponded to the relative mRNA levels except for GluN2A. The number of GluN2A puncta (10 puncta / 25 µm^2^, *SD* 2) was significantly lower than GluN2B (18 puncta / 25 µm^2^) and similar to GluN2C (9 puncta / 25 µm^2^; **Figure 3F**). The lower GluN2A protein levels might be due to weaker GluN2A antibody binding relative to other GluN2 antibodies or reduced synthesis/stability of GluN2A protein compared to other GluN2 subunits. GluN2 antibodies were validated for western blotting and immunohistochemistry on tissue from *Grin2a, Grin2c,* and *Grin2d* null mice as well as the cerebellum from 12-week old mice, which has very low levels of GluN2B and high GluN2A and GluN2C expression (Monyer et al., 1994)(**Supplemental Figures S3 – S5**). Altogether, these data demonstrate that all four GluN2 subunits exist at appreciable levels in the thalamus of 4-week-old mice, and that distinct transcriptional and/or post-transcriptional regulatory mechanisms likely control NMDAR subunit mRNA and protein levels.

**Figure 3.**
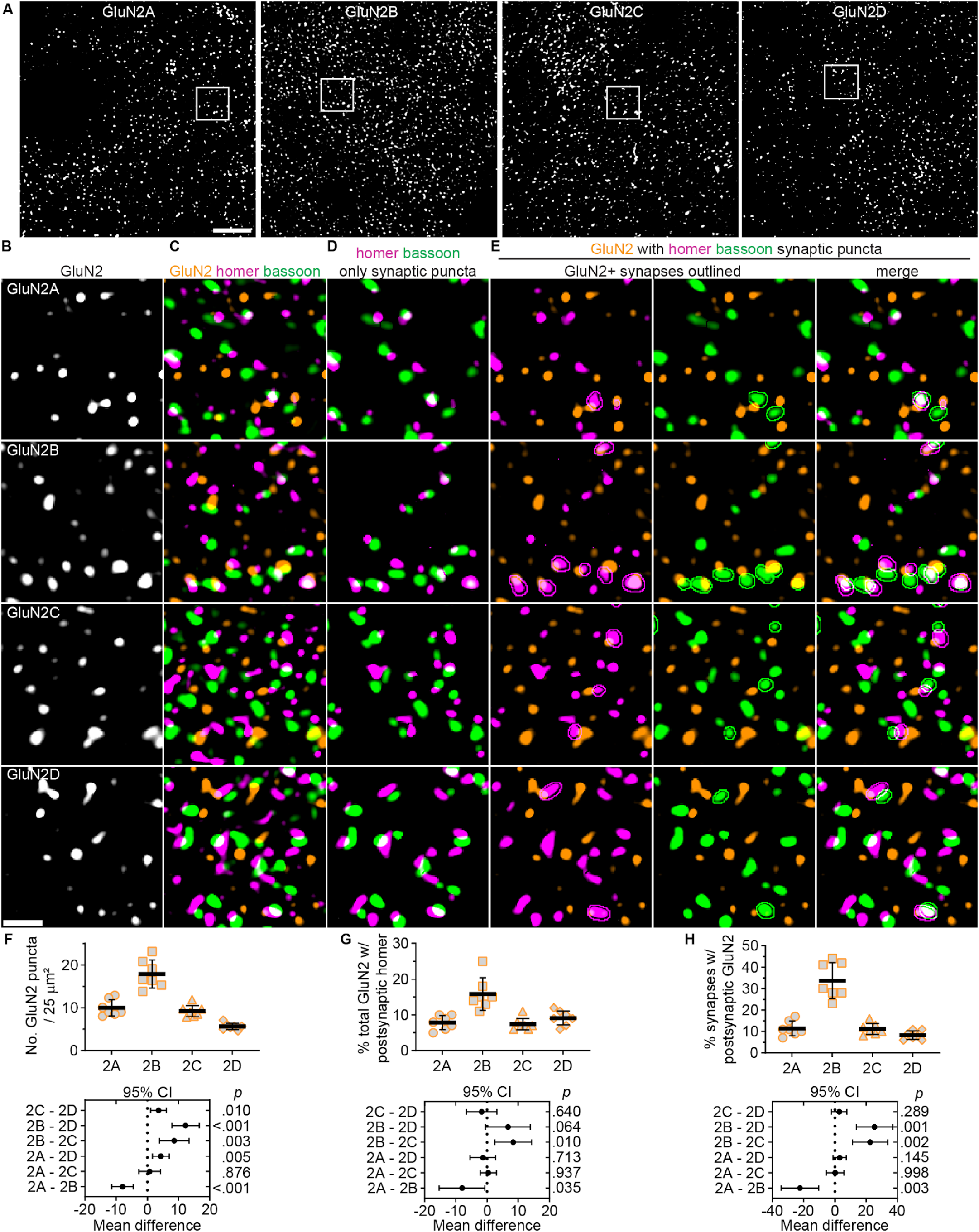
Synaptic localization of NMDARs differs among GluN2 subtypes. **A.** 60x (4x SoRa) images show GluN2A – GluN2D immunostaining in VPM thalamus. Boxed regions from panel A show (**B**) GluN2 signals, (**C**) merged images of GluN2, homer, and bassoon, (**D**) thresholded images of only homer/bassoon synaptic puncta (see **Supplemental Figure S7** for raw homer/bassoon images), and (**E**) merged images of GluN2 and synaptic homer or bassoon with outlines around GluN2-positive synapses identified by GluN2 overlapping with postsynaptic homer. **F**. GluN2 puncta number per 25 µm^2^ (the area of the boxed regions), **G**) the percentage of GluN2 overlapping postsynaptic homer, and **H**) the percentage postsynaptic homer overlapping GluN2 were quantified in the full FOV, averaged across three images per mouse, and plotted with group mean ± *SD* (n = 7 mice; 4F, 3M). All data were analyzed with a mixed effects model and Tukey’s pairwise comparisons for which the 95% CI of the mean difference are plotted beneath the group data with the *p* values on the right y-axis. **F:** *F* (2.01, 12.1) = 48.6, *p <* 0.001; **G**: *F* (1.80, 14.4) = 13.5, *p <* 0.001; **H**: *F* (1.52, 9.14) = 41.9, *p <* 0.001. Scale bars: (A) 20 µm, (C) 1 µm.

GluN2 subunit synaptic localization was assessed first with a NN analysis to quantify the correlation between the spatial distributions of homer with GluN2A – 2D (**Supplemental Figure S6**). This analysis served as an unbiased means for assessing whether all four GluN2 subunits were likely localized to homer/bassoon synapses. If any GluN2-homer interaction value was near zero, this would indicate the proteins were randomly distributed relative to one another; therefore, any synaptic localization observed in subsequent analyses would likely be random overlap. Homer localization was correlated with all GluN2 subunits, but showed a significantly weaker interaction strength with GluN2A (1.49, *SD* 0.76) compared to GluN2B (3.38, *SD* 1.4). Interaction strengths for GluN2C (2.65, *SD* 1.18) and GluN2D (2.0, *SD* 1.13) were intermediate and not significantly different from either GluN2A or GluN2B. These data suggest that all GluN2 subunits are likely localized to homer/bassoon synapses, but proximity to homer and/or abundance at synapses differ across GluN2 subtypes.

To directly assess GluN2 synaptic localization, we quantified the percentage of GluN2 puncta localized to homer/bassoon synapses as well as the percentage of homer/bassoon synapses overlapping with each GluN2 subunit (**Figure 3B – E** and **Supplemental Figure S7**). A larger percentage of GluN2B puncta (16%, *SD* 5) overlapped postsynaptic homer compared to GluN2A (8%, *SD* 2), GluN2C (7%, *SD* 2), and GluN2D (9%, *SD* 2; **Figure 3G**). This suggests that GluN2B is localized to synapses most efficiently. Consistent with this, the percentage of homer/bassoon synapses with postsynaptic GluN2B (34%, *SD* 8) was significantly higher than GluN2A (11%, *SD* 3), GluN2C (11%, *SD* 3), and GluN2D (8%, *SD* 2; **Figure 3H**), which were similar to one another. Likewise, the percentage of homer/bassoon synapses with presynaptic GluN2B (9%, *SD* 2) was significantly higher than GluN2A (5%, *SD* 1), GluN2C (5%, *SD* 1), and GluN2D (5%, *SD* 1; **Supplemental Figure S8**). Altogether, these data suggest that the majority of synaptic GluN2 subunits are postsynaptic with limited presynaptic localization, and GluN2B is the most abundant NMDAR subunit at glutamatergic VPM synapses.

Given that GluN2B had the highest expression level, we expected that it would also be the most abundant subunit at synapses. However, based on their mRNA and proteins levels, we predicted GluN2A and GluN2C would be expressed at a greater proportion of synapses than observed. We surmised that, in addition to being localized to synapses less frequently than GluN2B, the GluN2A- and GluN2C-containing NMDARs present at synapses may be localized farther from the postsynaptic density compared to GluN2B. To assess this, we compared the frequency distribution of distances between postsynaptic GluN2 subunits and postsynaptic homer (**Supplemental Figure S9**). The median distance was significantly shorter for GluN2B (153 nm, *IQR* 24) compared to GluN2A (193 nm, *IQR* 36), GluN2C (196 nm, *IQR* 41), and GluN2D (204, *IQR* 52). These data suggest that GluN2B-containing NMDARs were localized closest to the postsynaptic density at VPM synapses.

### Preferential localization of GluN2 subunits at CT and sensory synapses in the VPM

We hypothesized that the observed differences in GluN2 synaptic localization and distance from postsynaptic homer could have been due to input-specific organization of GluN2 subtypes at CT and sensory synapses. To address this hypothesis, we identified CT and sensory inputs by detecting homer/bassoon synapses and simultaneously immunostaining for VGLUT1 or VGLUT2, which labeled CT and sensory axon terminals, respectively (**Figure 4A,B**). Consistent with previous reports, sensory synapses were few in number and formed by large presynaptic terminals with multiple release sites, whereas CT terminals were numerous and small. Synapses containing VGLUT1-positive bassoon puncta comprised 49% (*SD* 10) of all detected homer/bassoon synapses, and 6% (*SD* 2) of synapses had VGLUT2-positive bassoon puncta (**Figure 4C – E**). NMDARs contribute to postsynaptic currents at both CT and sensory synapses and, thus, were expected to be detectable at both synapse populations. We immunolabeled VGLUT1 or VGLUT2 along with homer, bassoon, and GluN1 (**Figure 4F,G**) and detected postsynaptic GluN1 at 53% (*SD* 8) of VGLUT1-positive synapses and 45% (*SD* 11) of VGLUT2-positive synapses (**Figure 4H**). Presynaptic GluN1 was detected at 8% (*SD* 2) of VGLUT1-positive synapses and 11% (*SD* 3) of VGLUT2-positive synapses (**Figure 4I**). These data demonstrate that we can specifically label CT and sensory synapses and detect NMDARs at similar proportions of both synapse types.

**Figure 4.**
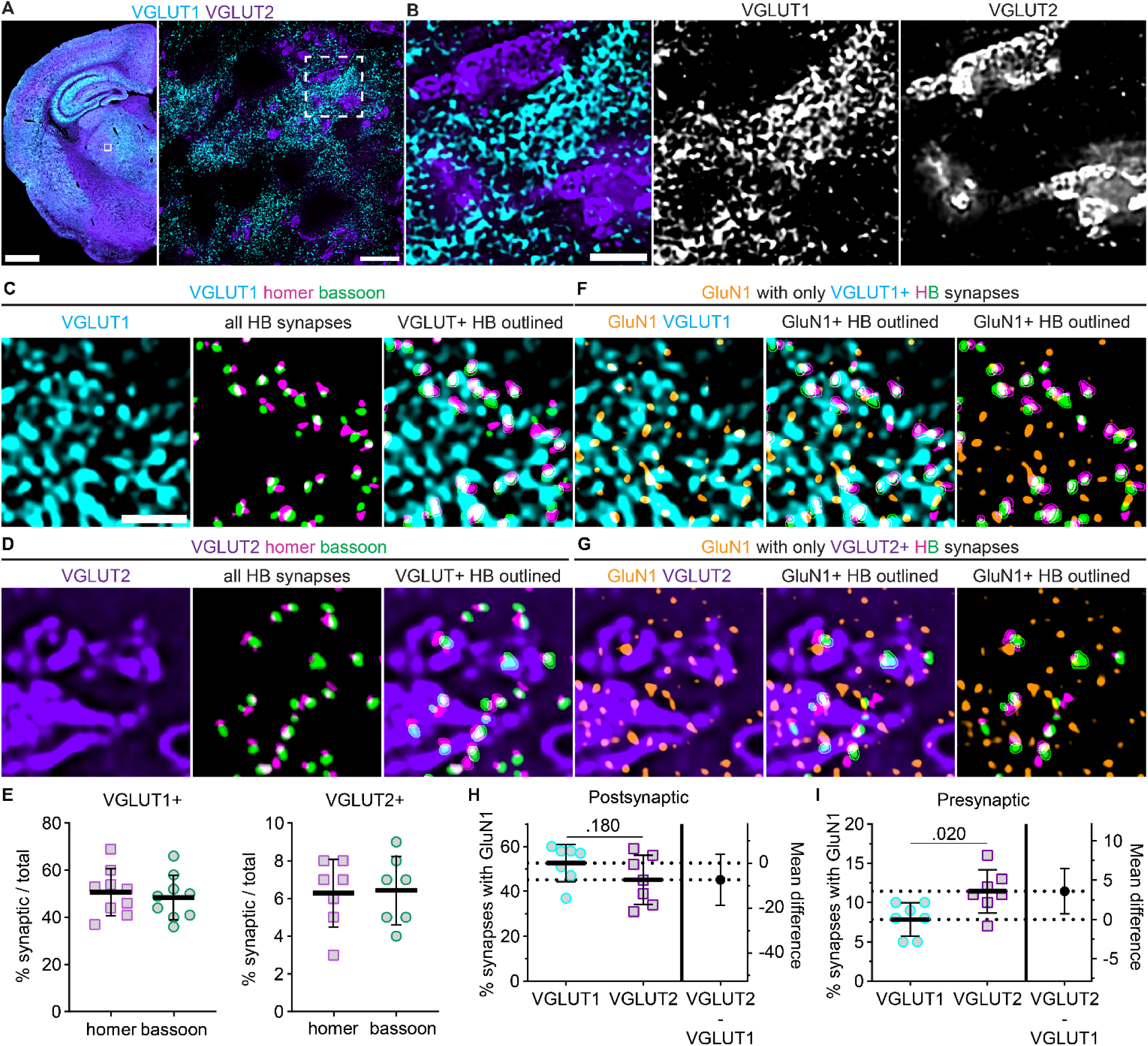
NMDAR localization at VGLUT1- and VGLUT2-positive synapses. **A.** VGLUT1 and VGLUT2 immunostaining is shown in a tiled 10x image of a mouse brain section and 60x (4x SoRa) images of the boxed region. **B.** The dashed region from panel A is expanded to visualize non-overlapping VGLUT1- and VGLUT2-positive synaptic terminals. 60x (4x SoRa) images show (**C**) VGLUT1 or (**D**) VGLUT2 immunostaining alongside thresholded signals of all homer/bassoon (HB) synaptic puncta, and merged images of VGLUT1/2 with VGLUT-positive HB synapses outlined. **E**. The percentage of total synapses with bassoon puncta overlapping VGLUT1 (n = 9 mice; 4F, 5M) and VGLUT2 (n = 7 mice; 4F, 3M) were plotted with the group mean ± *SD*. **F,G.** 60x (4x SoRa) images show GluN1 and VGLUT1/2 immunostaining (left) with GluN1 and thresholded signals of only VGLUT-positive synapses with GluN1-positive synaptic puncta outlined (middle and right). The percentage of VGLUT-positive synapses with GluN1 overlapping (**H**) homer (postsynaptic) and (**I**) bassoon (presynaptic) were plotted (n = 7 mice; 4F, 3M) with the mean ± *SD*. Groups were compared with Welch’s t tests (**H**: *t =* 1.433, df = 11.16; **I**: *t* = 2.717, df = 11.24). Scale bars: (**A**) 1 mm, 20 µm; (**B**) 3 µm; (**C**) 1 µm.

To examine input-specific GluN2 localization in the VPM thalamus, we quantified the fraction of VGLUT1- and VGLUT2-positive homer puncta overlapping GluN2 (**Figure 5A,B**). We were unable to achieve efficient four channel labeling for all GluN2 subunits with homer, bassoon, and VGLUT1 or VGLUT2 with the available antibodies. We surmised that at most synapses detected in our images both bassoon and homer would be overlapping the VGLUT signal to some extent. We utilized those subunit combinations that allowed four channel immunolabeling to test whether quantifying the percentage of VGLUT-positive homer/bassoon synapses with homer overlapping GluN2 (as done in Figure 4 with GluN1) yielded similar results to quantifying the percentage of VGLUT-positive homer puncta overlapping GluN2 without bassoon labeling (**Supplementary Figure S10**). While the total number of homer puncta was greater than the total number of bassoon/homer synapses detected, the relative percentage of VGLUT1- and VGLUT2-positive homer overlapping GluN2 was not significantly different. Therefore, we assessed input-specific GluN2 localization by quantifying the percentage of VGLUT1- or VGLUT2-positive homer puncta that overlapped each GluN2 subunit.

**Figure 5.**
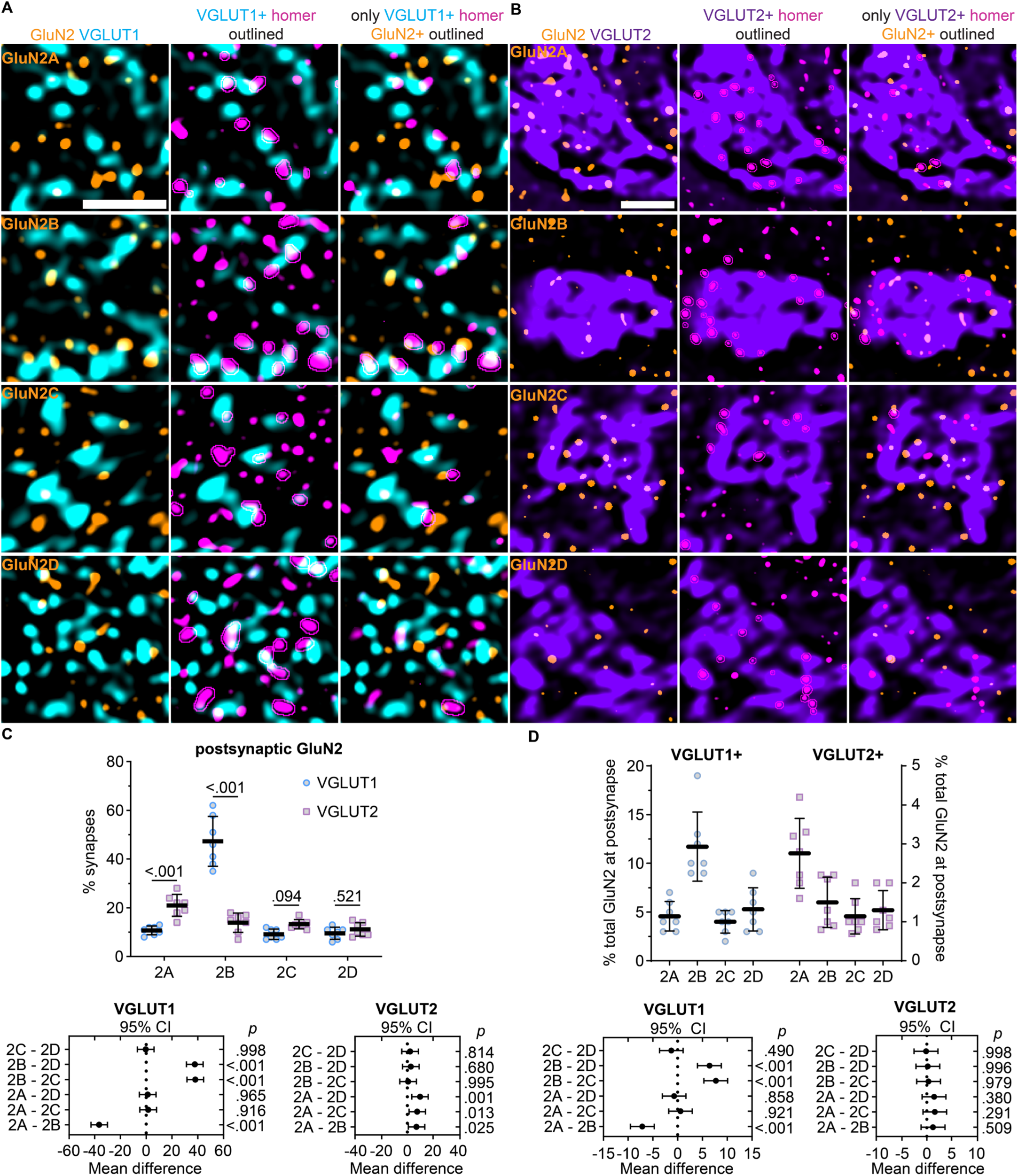
GluN2A and GluN2B are preferentially localized to different VPM inputs. **A.** 60x (4x SoRa) images show GluN2A – 2D immunostaining with VGLUT1 (left); VGLUT1 and homer with VGLUT1-positive homer outlined (middle), and GluN2, VGLUT1, and only VGLUT1-positive homer with GluN2-positive homer puncta outlined (right). **B**. 60x (4x SoRa) images show GluN2A – 2D immunostaining with VGLUT2 (left); VGLUT2, and VGLUT2-positive homer outlined (middle); and GluN2, VGLUT2, and only VGLUT2-positive homer with GluN2-positive homer puncta outlined (right). **C**. The percentage of VGLUT1- and VGLUT2-positive homer puncta overlapping GluN2 (n = 7; 4F, 3M) were plotted with the group mean ± *SD*. The data were compared by a mixed effects model with Tukey’s pairwise comparisons [F(3,48) = 66.0; p < 0.001]. The p values for input-specific comparisons with each GluN2 subunit are shown on the plot. The 95% CI of the mean difference for comparisons across GluN2 subunits within each input type are plotted beneath the group data with *p* values on the right y-axis. **D**. The percentage of each GluN2 subunit overlapping VGLUT1/2-positive homer puncta were plotted and compared by mixed effects model with Tukey’s pairwise comparisons [F (3, 18) = 22.6, p < 0.001]. The 95% CI of the mean difference are plotted beneath the group data with *p* values on the right y-axis. Scale bars: (**A**) 1 µm; (**B**) 5 µm.

GluN2B overlapped with 47% (*SD* 10) of VGLUT1-positive homer puncta, but only 13% (*SD* 4) of VGLUT2-positive homer puncta (**Figure 5C**). On the contrary, GluN2A overlapped with a significantly greater proportion of VGLUT2-positive homer puncta (21%, *SD* 5) than VGLUT1-positive homer puncta (11%, *SD* 2). GluN2C and GluN2D overlapped with similar percentages of VGLUT1-positive (2C: 9%, *SD* 2; 2D: 10%, *SD* 2) and VGLUT2-positive homer puncta (2C: 13%, *SD* 2; 2D: 11%, *SD* 3). These data suggest that GluN2B is the most abundant subunit at CT synapses, whereas GluN2A is the most abundant subunit at sensory synapses. Notably, these results suggest that GluN2C and GluN2D are expressed at a similar proportion of sensory synapses as GluN2B, despite having substantially lower expression levels. To determine how efficiently GluN2A – 2D were localized to CT and sensory synapses, we quantified the percentage of total GluN2A – 2D puncta that were overlapping VGLUT1- and VGLUT2-positive homer. This analysis demonstrated that 12% (*SD* 4) of total GluN2B puncta overlapped with VGLUT1-positive homer, which was significantly more than GluN2A (5%, *SD* 2), GluN2C, (4%, *SD* 1), and GluN2D (5%, *SD* 2; **Figure 5D**). In contrast, 3% (*SD* 1) of total GluN2A overlapped with VGLUT2-positive homer, while only 1% (*SD* 0.5) of total GluN2B, GluN2C, and GluN2D overlapped with VGLUT2-positive homer. Given that approximately 90% of glutamatergic synapses detected in the VPM were VGLUT1-positive and only 10% were VGLUT2-positive, these data suggest that GluN2A and GluN2B were preferentially localized to sensory and CT synapses, respectively.

### NMDAR distance from the synapse is input- and GluN2 subunit-specific

Since GluN2B was preferentially localized to VGLUT1-positive synapses and had the shortest distance to postsynaptic homer when analyzing all VPM synapses, we reasoned that NMDARs at VGLUT1-positive synapses may be localized closer to homer than those at VGLUT2-positive synapses, where GluN2A is overrepresented. On the other hand, the distance from NMDARs to homer could be determined primarily by GluN2 subunit composition and not by synapse identity. To address this question, we measured the distances between NMDAR subunits and homer at VGLUT1- and VGLUT2-positive synapses, plotted the data as frequency distributions, and compared the median distances (**Supplemental Figure S11A,B**). We first assessed all NMDARs by comparing GluN1-homer distances and found that the median distance for NMDARs at VGLUT1-positive synapses (147 nm, *SD* 11) was shorter than VGLUT2-positive synapses (177 nm, *SD* 18; **Supplemental Figure S11C**). These data suggest that NMDARs at VGLUT1-positive synapses are localized closer to the postsynaptic density than VGLUT2-positive synapses. Analyzing GluN2-homer distances revealed that GluN2B and GluN2C distances from homer were input-specific (**Supplemental Figure S11D**). GluN2B was localized significantly closer to VGLUT1-positive homer puncta (163 nm, *SD* 19) compared to VGLUT2-positive homer (189 nm, *SD* 27). GluN2C was localized significantly closer to VGLUT2-positive homer (177 nm, *SD* 19) compared to VGLUT1-positive synapses (198 nm, *SD* 14). GluN2A and GluN2D were localized similar distances from VGLUT1- (2A: 207 nm, *SD* 15; 2D: 204 nm, *SD* 21) and VGLUT2- (2A: 190 nm, *SD* 10; 2D: 185 nm, *SD* 22) positive homer puncta. These data suggest that NMDAR subsynaptic organization is both GluN2 subunit-specific and input-specific in VPM thalamus.

### GluN2 subtypes show preferential colocalization with synaptic scaffolding proteins

We hypothesized that different postsynaptic scaffolding proteins might be governing the input-specific organization of GluN2 subunits at VPM synapses. To begin to investigate this, we performed a NN analysis to correlate the spatial distributions of all NMDARs relative to PSD95 and SAP102, two scaffolding proteins known to interact with NMDARs (**Supplemental Figure S12A - D**)(Gardoni & Di Luca, 2021). The NN interaction strength for GluN1 with SAP102 (2.25, *SD* 0.38) was significantly higher than PSD95 (1.93, *SD* 0.48) and similar to GluN1/homer interaction strength (2.84, *SD* 0.66). Furthermore, GluN1 was localized to a greater fraction of synapses detected by SAP102/bassoon pairs (49%, *SD* 6) compared to PSD95/bassoon pairs (28%, *SD* 6) and a similar fraction of synapses as detected by homer/bassoon pairs (49%, *SD* 12). Notably, homer, PSD95, and SAP102 labeling showed no significant difference in NN interaction strength with bassoon, number of synapses detected, and percentage of total bassoon detected as presynaptic (**Supplemental Figure S12E – J**), indicating that all three pairs are effective markers of synapses in the thalamus. These data suggest that NMDARs as a whole show preferential colocalization with SAP102- and homer-positive synapses compared to PSD95-positive synapses in the VPM thalamus.

Further NN analyses showed that the spatial distributions of all GluN2 subunits were significantly correlated with PSD95 and SAP102, but the interaction strengths differed across GluN2 subunits (**Figure 6A - C)**. PSD95 localization was weakly correlated with GluN2A (1.80, *SD* 0.29) and GluN2D (2.20, *SD* 0.90), but was highly correlated with GluN2B (4.76, *SD* 0.92) and GluN2C (3.98, *SD* 0.81). On the other hand, SAP102 localization was associated with GluN2A (5.40, *SD* 1.30), GluN2B (4.49, *SD* 0.65), GluN2C (2.49, *SD* 0.88), and GluN2D (3.73, *SD* 0.61). These data showed that scaffolding proteins preferentially localized with specific GluN2 subunits. We also determined that GluN2 subunits preferentially localized with specific postsynaptic scaffolding proteins by comparing the NN interaction strengths for each GluN2 subunit across PSD95, SAP102, and homer. GluN2A NN interaction strength with SAP102 was significantly greater than PSD95 and homer (1.32, *SD* 0.5). GluN2C-PSD95 interaction strength was significantly greater than SAP102 and homer (2.39, *SD* 0.8). GluN2B interaction strength was similar for PSD95, SAP102, and homer (3.27, *SD* 0.9), while GluN2D was most strongly associated with SAP102 compared to PSD95 and homer (2.54, *SD* 1.1). These data suggest that GluN2 subunits colocalize with different synaptic proteins, which could contribute to their input-specific organization. This observation further suggested that assessing GluN2 synaptic localization with homer/bassoon puncta may have underestimated the synaptic localization of GluN2A, GluN2C, and GluN2D, which had a higher NN interaction with SAP102 and/or PSD95 compared to homer.

**Figure 6.**
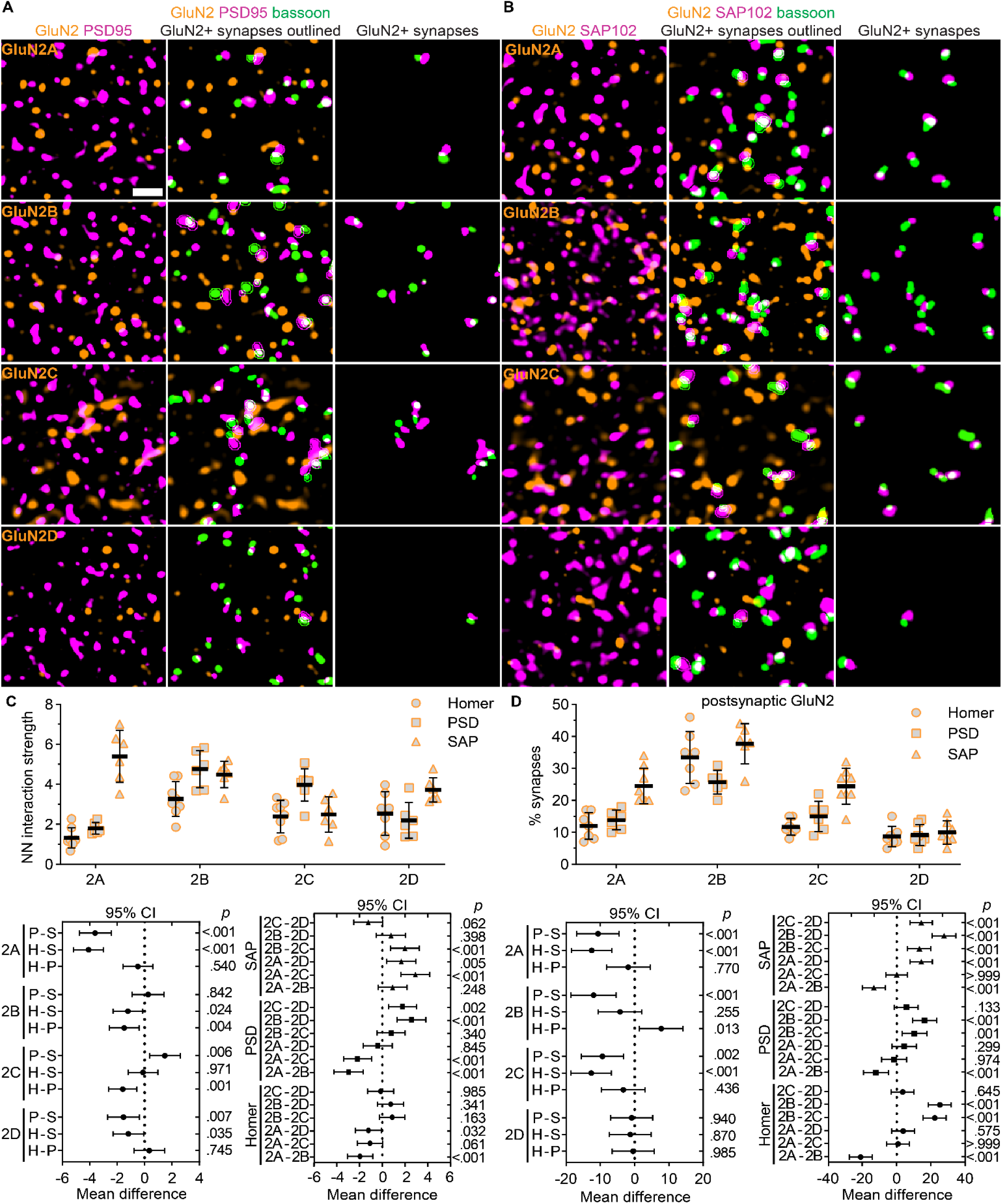
Preferential colocalization of GluN2 subunits with synaptic scaffolding proteins. **A.** 60x (4x SoRa) images show GluN2A – 2D immunostaining with PSD95 (left); PSD95 and bassoon with GluN2-positive synaptic puncta outlined (middle), and only GluN2-positive synapses (right). **B**. 60x (4x SoRa) images show GluN2A – 2D immunostaining with SAP102 (left); SAP102 and bassoon with GluN2-positive synaptic puncta outlined (middle), and only GluN2-positive synapses (right). **C**. The NN interaction strength for GluN2 with homer, PSD95, and SAP102 (n = 7; 3F, 4M) were plotted with the group mean ± *SD*. The data were compared by a two-way ANOVA with Tukey’s pairwise comparisons [F (6, 68) = 12.8, *p* < 0.001] with the 95% CI of the mean difference plotted beneath the group data for comparisons across postsynaptic markers for each GluN2 subunit (bottom, left) and across GluN2 subunits for each postsynaptic marker (bottom, right). The *p* values are on the right y-axis. **D.** The percentage of synapses with postsynaptic puncta overlapping GluN2 (n = 7; 3F, 4M) were plotted with the group mean ± *SD*. The data were compared by a mixed effects model with Tukey’s pairwise comparisons [F (6,70) = 4.22, *p* = 0.001] with the 95% CI of the mean difference plotted beneath the group data for comparisons across postsynaptic markers for each GluN2 subunit (bottom, left) and across GluN2 subunits for each postsynaptic marker (bottom, right). The *p* values are on the right y-axis. Scale bar: (**A**) 1 µm.

To address this, we detected synapses with bassoon/homer, bassoon/PSD95, or bassoon/SAP102 pairs and quantified the proportion of synapses with the postsynaptic marker overlapping each GluN2 subunit (**Figure 6D**). The percentages of postsynaptic homer puncta overlapping with GluN2A (12%, *SD* 4), GluN2B (33%, *SD* 8), GluN2C (12%, *SD* 3), and GluN2D (9%, *SD* 3) were consistent with Figure 3F. The percentage of postsynaptic PSD95 puncta overlapping with GluN2A (14%, *SD* 3), GluN2C (15%, *SD* 5), and GluN2D (9%, *SD* 3) were similar to homer, but the percentage overlapping with GluN2B (26%, *SD* 4) was significantly reduced compared to homer. Most notably, the percentage of postsynaptic SAP102 overlapping with GluN2A (25%, *SD* 6) and GluN2C (24%, *SD* 6) were significantly higher than for postsynaptic homer and PSD95. Postsynaptic SAP102 overlap with GluN2B (37%, *SD* 6) and GluN2D (10%, *SD* 4) were similar to homer. These data suggest that GluN2A and GluN2C are localized to a greater fraction of synapses than initially detected utilizing homer/bassoon labeling and may be preferentially localized to SAP102-containing synaptic domains.

### GluN2 localization to sensory synapses differs based on postsynaptic marker

Given that GluN2A showed preferential localization to VGLUT2-positive synapses and SAP102-containing synapses, we surmised that SAP102 may be a better marker of VGLUT2-positive synapses than homer and, thus, would provide a better means for assessing postsynaptic NMDAR localization at sensory synapses. To test this, we compared the percentage of total bassoon puncta that were identified as part of VGLUT2-positive synapses when combined with homer and SAP102 (**Figure 7A**). VGLUT2-positive synapses detected by SAP102/bassoon included 11% (*SD* 3) of total bassoon puncta while homer/bassoon pairs included only 6% (*SD* 2; **Figure 7B**). Furthermore, a greater percentage of total SAP102 puncta (11%, *SD* 3) were detected as part of VGLUT2-positive synapses compared to homer (6%, *SD* 2). Consistent with these findings, a greater percentage of total VGLUT2-positive bassoon puncta were identified as presynaptic puncta when combined with SAP102 (38%, *SD* 6) compared to homer (26%, *SD* 4; **Figure 7C**). Given that we found no significant differences for the NN interaction strengths or the total number of synapses detected by homer/bassoon and SAP102/bassoon (**Supplemental Figure S11**), these data suggest that SAP102/bassoon labeling detected VGLUT2-positive synapses more effectively homer/bassoon labeling.

**Figure 7.**
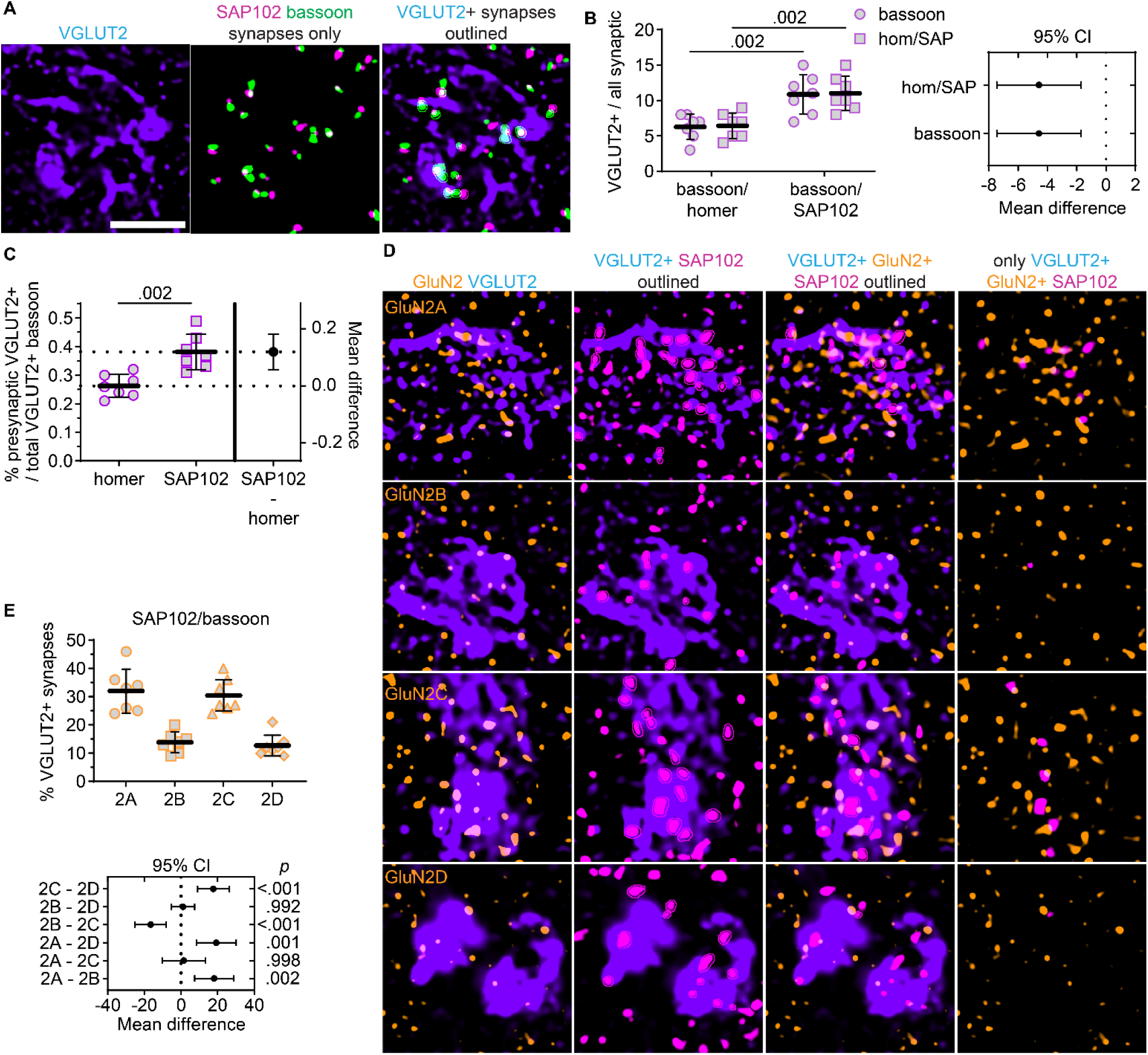
GluN2A and GluN2C are preferentially localized with VGLUT2-positive SAP102. **A.** 60x (4x SoRa) images show VGLUT2 immunostaining (left), a thresholded image depicting SAP102 and bassoon puncta detected as synapses (middle), and a merged image with VGLUT2-positive synaptic puncta outlined. **B.** The percentage of total bassoon puncta (circles) and homer or SAP102 puncta (squares) that were detected as VGLUT2-positive synapses were plotted with the group mean ± *SD* (n = 7; 4F, 3M). The data were compared by a mixed effects model [F (1, 12) = 15.7, *p* = 0.002] with Sidak’s comparisons. The 95% CI of the mean difference for comparisons of bassoon and homer/SAP102 are plotted (right). **C**. The percentage of total VGLUT2-positive bassoon that were detected as part of VGLUT2-positive synapses are shown in an estimation plot with group mean ± *SD*. The data were compared by a Welch’s t-test with the 95% CI of the mean difference plotted on the right y-axis (*t* = 4.27, *df* = 10.2). **D**. 60x (4x SoRa) images show GluN2A – 2D immunostaining with VGLUT2 (left); VGLUT2 and SAP102 with postsynaptic VGLUT2-positive SAP102 outlined; GluN2, VGLUT2, and only VGLUT2-positive SAP102 with GluN2-positive SAP102 puncta outlined; and GluN2 with only VGLUT2- and GluN2-positive SAP102 puncta. **E**. The percentage of VGLUT2-positive SAP102 puncta overlapping GluN2 were plotted with the group mean ± *SD*. The data were compared by a mixed effects model with Tukey’s pairwise comparisons, and the 95% CI of the mean difference for comparisons across GluN2 subunits are plotted beneath the group data with *p* values on the right y-axis [F (1.68, 13.5) = 23.9, *p* < 0.001]. Scale bars (**A**) 2 µm.

Based on this finding, we compared GluN2 localization at VGLUT2-positive SAP102 (**Figure 7D**). The percentage of VGLUT2-positive SAP102 puncta overlapping GluN2A (32%, *SD* 8) and GluN2C (30%, *SD* 6) were significantly greater than GluN2B (14%, *SD* 4) and GluN2D (13%, *SD* 4; **Figure 7E**). These data indicate that both GluN2A and GluN2C are preferentially localized with VGLUT2-positive SAP102. This GluN2A result is consistent with our findings at homer/bassoon synapses, but SAP102 synapse detection revealed greater GluN2C localization to VGLUT2-positive synapses than homer. Altogether, these data indicate that GluN2 subunits and postsynaptic scaffolding proteins exhibit input-specific synaptic organization in the VPM thalamus.

## DISCUSSION

NMDAR diversity contributes to synapse heterogeneity across and within brain regions, but few regions allow for directly comparing the localization and function of diverse NMDAR subtypes in a single neuron population. We took advantage of the extensive NMDAR diversity in the thalamus to determine how all four GluN2 subtypes were organized at two anatomically distinct glutamatergic inputs. We discovered subtype- and input-specific NMDAR synaptic localization and subsynaptic organization in VPM thalamocortical neurons. GluN2 subunits were also preferentially localized with specific postsynaptic scaffolding proteins, which provides insight into potential mechanisms underlying input-specific localization of NMDAR subtypes. Beyond uncovering molecular diversity among thalamic synapses, this study highlights the utility of thalamocortical neurons and super resolution imaging for investigating the molecular mechanisms that generate glutamatergic synapse diversity in native thalamus.

CT and sensory synapses in the thalamus exhibit distinct physiological properties that impact their development, basal neurotransmission, and plasticity (Golshani et al., 1998; Hsu et al., 2010, 2012). Input-specific GluN2B function likely contributes to these differences as GluN2B-containing NMDARs mediate synaptic transmission at both CT and sensory synapses in early development, but only CT synapses late in postnatal development and adulthood (Miyata & Imoto, 2006). Sensory synapses have a much larger AMPAR/NMDAR ratio than CT synapses late in postnatal development, but a substantial NMDAR current remains. Physiological studies have not identified which NMDAR subtypes contribute to this sensory synaptic current. Consistent with these earlier findings, we observed GluN2B localization to a large proportion of CT synapses, but only a small proportion of sensory synapses in 4-week-old mice. On the other hand, GluN2A and GluN2C were localized to a large proportion of sensory synapses and a small proportion of CT synapses. These data suggest that, as GluN2A and GluN2C expression increase from 2 – 4 weeks of age, these subunits might replace GluN2B at sensory synapses, but not at CT synapses. Similar developmental increases have been established for GluN2A in the cortex and hippocampus, and for GluN2A and GluN2C in cerebellar granule cells (Ewald & Cline, 2009). The unique input-specific nature of the developmental shift in VPM thalamus suggests TC neurons have both temporal and spatial regulatory mechanisms controlling NMDAR localization in a GluN2-specific manner.

In addition to differential synaptic localization of GluN2 subunits, we observed that the distance between postsynaptic markers and GluN2 subunits was input- and subtype-specific. In general, NMDARs at CT synapses were localized closer to the postsynaptic density (labeled by homer) than those at sensory synapses. NMDAR localization farther from the postsynaptic density could contribute to the reduced NMDAR current at sensory synapses. NMDARs farther from the synapse could be in an intracellular reserve pool or activated by glutamate spillover, perhaps only during high frequency stimulation. In addition, high GluN2C expression at sensory synapses would also yield smaller NMDAR currents than a similar number of GluN2B-containing NMDARs at CT synapses due to their distinct biophysical properties. Large proportions of GluN2A, GluN2C, and GluN2D were not localized to synapses in general, suggesting that GluN2B might be trafficked more efficiently to synapses. Interestingly, higher order TC neurons express greater levels of GluN2C and GluN2D relative to GluN2A and GluN2B than sensory TC neurons (Phillips et al., 2019). Given that higher order thalamic nuclei do not have sensory inputs, it is possible that GluN2A, GluN2C, and GluN2D expression are higher at CT inputs relative to sensory nuclei. Additionally, higher order nuclei receive inputs from disparate cortical areas, and perhaps GluN2 expression is preferentially localized among anatomically distinct cortical inputs to these cells. Some reticular thalamus neurons and higher order TC neurons also express GluN3A, for which virtually no information exists in terms of synaptic localization or function in the thalamus. It will be intriguing to determine how the synaptic localization of NMDAR subunits, and their respective physiological roles, differ in higher order TC neurons compared to sensory TC neurons.

While we have elucidated the input-specific organization of all four GluN2 subunits, their contributions to synapse function remain unclear in TC neurons. The functional properties of GluN2 subunits are vastly different, but how it is advantageous to have specific functionalities at CT and sensory synapses remains unclear. We can speculate that perhaps the fast decay of GluN2A-containing NMDARs could be important at sensory synapses, which have predominant AMPAR-mediated currents and where the rate of synaptic input carries essential information about somatosensation and pain (Kenshalo et al., 1980; Yen & Lu, 2013). Furthermore, plasticity has been observed at CT synapses, but not sensory synapses (Castro-Alamancos, 2002; Castro-Alamancos & Calcagnotto, 1999). In the hippocampus, GluN2B has been identified as promoting long-term synaptic plasticity, whereas GluN2A expression may restrict plasticity; although, this is not universal in the brain and can be context-dependent (Shipton & Paulsen, 2014). Nevertheless, GluN2B localization to CT synapses and sensory synapses early in development could enable synaptic plasticity, while the removal of GluN2B from sensory synapses later in development could serve to restrict plasticity of sensory pathways. The functional contributions of GluN2A, GluN2C, and GluN2D are poorly understood at TC neuron synapses as are their roles in thalamic physiology more generally.

One limitation of this study is that we did not distinguish between in diheteromeric or triheteromeric NMDARs. It is possible that GluN2A and GluN2C expression at sensory synapses exist primarily as triheteromeric receptors similar to the cerebellum (Bhattacharya et al., 2018). Given that GluN2A and GluN2B were localized preferentially to distinct inputs, GluN1/2A/2B triheteromers may not be as prevalent in thalamus as in cortex and hippocampus (Al-Hallaq et al., 2007). We utilized the SoRa spinning disk method as this imaging modality allowed us to efficiently label and visualize numerous combinations of NMDARs and synaptic markers in native tissue. Co-labeling of multiple GluN2 subunits with this method could provide further insight as to how often these receptors are found close enough to be in the same tetramer. In addition, our analyses of single z-planes underestimated the true fraction of synapses expressing NMDARs, but this did not hinder our study’s objective of determining the *relative* efficiency of synaptic localization and subsynaptic organization among GluN2 subunits. Three-dimensional imaging would provide a more accurate view of how many synapses express each NMDAR subtype. Single-molecule imaging would also allow for more accurate distance measurements to determine NMDAR subsynaptic organization and provide a better assessment of how many NMDARs of each subtype are present at a single CT synapse or juxtaposed to a single release site at sensory synapses.

GluN2A and GluN2B subunits preferentially interact with particular synaptic protein complexes, and interactions with specific scaffolding proteins regulate their subsynaptic organization (Papouin & Oliet, 2014). For example, in the cortex, PSD95 and SAP102 were shown to be highly colocalized with GluN2A and GluN2B, respectively. Increasing PSD95 levels during development have been proposed to stabilize GluN2A at the synapse; whereas, SAP102 and GluN2B, which are more highly expressed early in development, exhibit greater mobility and move to extrasynaptic domains later in development. In the thalamus, synaptic scaffolding proteins showed preferential colocalization with particular GluN2 subunits and specific inputs suggesting these proteins could be involved in input-specific NMDAR localization. Notably, SAP102 preferentially colocalized with GluN2A at VGLUT2-positive sensory synapses, while the correlation between GluN2A and PSD95 localization was weaker. GluN2B was localized to a greater proportion of PSD95 synapses than GluN2A, and GluN2B was localized closer to the postsynaptic density than GluN2A. The distinct macromolecular complexes in which GluN2 subunits reside in the native thalamus are unknown, and this information remains largely unknown across the entire brain for GluN2C and GluN2D. However, our results thus far suggest that associations between NMDARs and scaffolding proteins as well as their synaptic organization may differ between the cortex and thalamus.

By uncovering synaptic NMDAR organization in the thalamus, this work establishes a tractable system for probing the mechanisms that regulate how specific subtypes are localized as well as how experience and disease may affect NMDAR localization in native brain tissue. Moreover, this study provides impetus for testing how NMDAR diversity contributes to the unique physiology of anatomically distinct synaptic inputs in the thalamus. Revealing GluN2-specific physiological roles is necessary to understand how it is advantageous for thalamocortical neurons to express such great diversity of NMDAR compared to pyramidal neurons in the cortex and hippocampus, for example. Uncovering the mechanisms by which GluN2 subunits and their associated macromolecular complexes direct input-specific localization will also provide insight into how the molecular signatures of synapses change across development. More broadly, investigating how these molecular signatures and regulatory mechanisms differ across brain circuits will advance our understanding of the molecular underpinnings of functional synapse diversity and circuit-specific vulnerability to disease.

## AUTHOR CONTRIBUTIONS

M.T., J.M., A.P-A., G.W. – data acquisition, manuscript editing; B.L.G. – experimental design, data acquisition, data analysis and interpretation, manuscript writing/editing; R.K. – data analysis, manuscript editing; S.F. – experimental design, data interpretation, and manuscript writing/editing; S.A.S. – project design and supervision, experimental design, data acquisition, data analysis and interpretation, funding acquisition, manuscript writing/editing.

## ACKNOWLEDGEMENTS

We would like to thank: Dr. Stephen Traynelis and Chad Camp for generously providing brain tissue from *Grin2a*, *Grin2c*, and *Grin2d* null mice; Christie Lacy and the Fralin Biomedical Research Institute at VTC Cellular Imaging Core facility for assistance with the Nikon SoRa spinning disk microscope.

