## Supplemental Figures for "Input-specific localization of NMDA receptor GluN2 subunits in thalamocortical neurons"

**Supplementary Information: Figures S1 – S12**

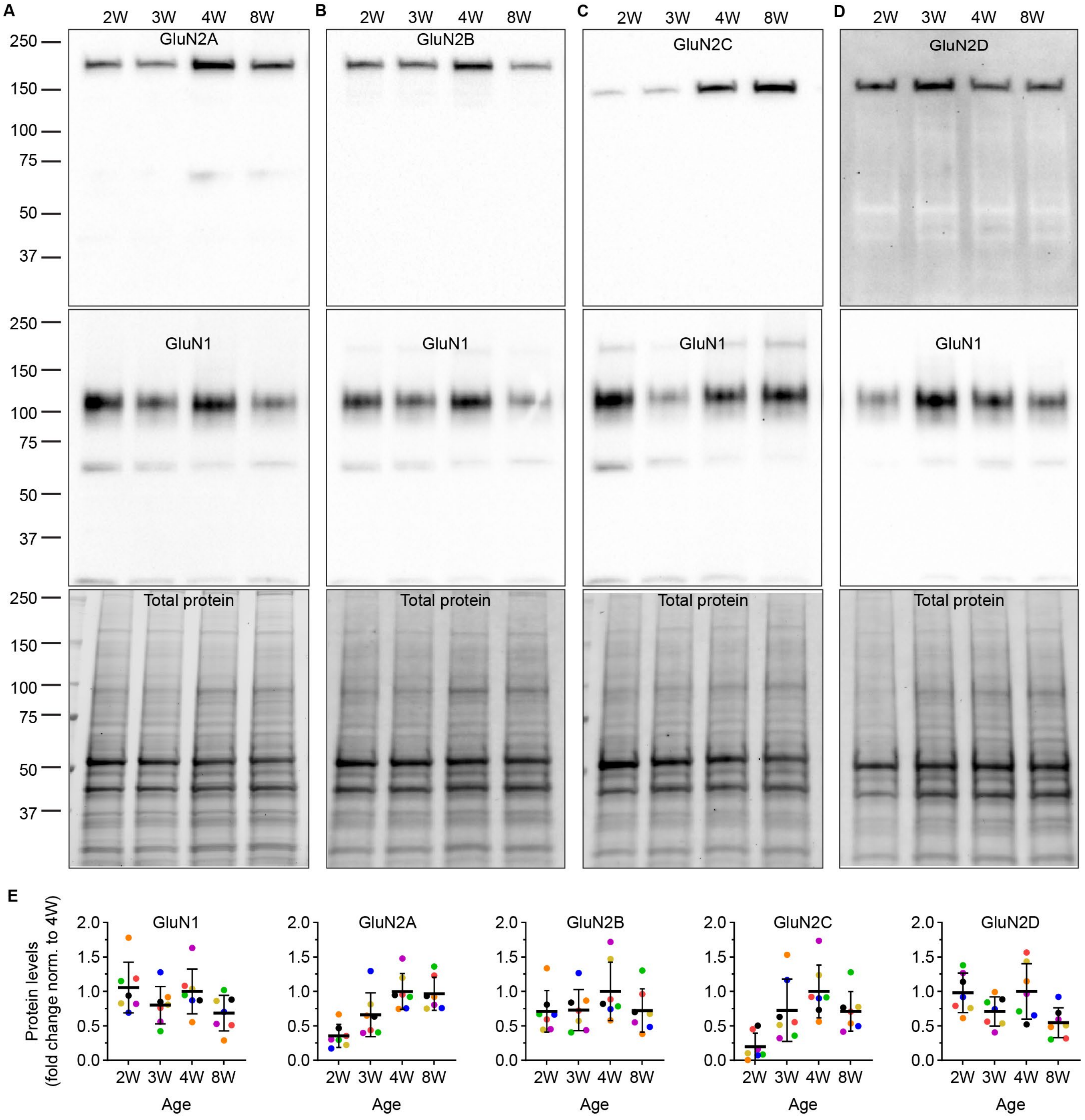


**Supplemental Figure S1. Developmental expression of GluN subunits in the thalamus.** VB thalamus tissue punches from C57Bl/6J mice aged 2, 3, 4, and 8 weeks (W) were homogenized, processed for SDS PAGE, run on duplicate gels, and then samples were immunoblotted for NMDAR subunits (*n* = 7 mice; 2W: sex unknown; 3W: 3F, 4M; 4W: 5M, 2F; 8W: 3M, 4F). Samples were probed for (**A**) GluN2A, (**B**) GluN2B, (**C**) GluN2C, and (**D**) GluN2D as well as GluN1 (middle, **A** – **D**). **E.** Protein levels were quantified by densitometry and normalized to total protein (bottom, **A** – **D**) within each gel. Protein levels for each replicate were shown as individual data points of the same color across age and expressed as fold change from the 4W mean to control for differences in band intensity across experiments. The line and error bars are mean ± SD.

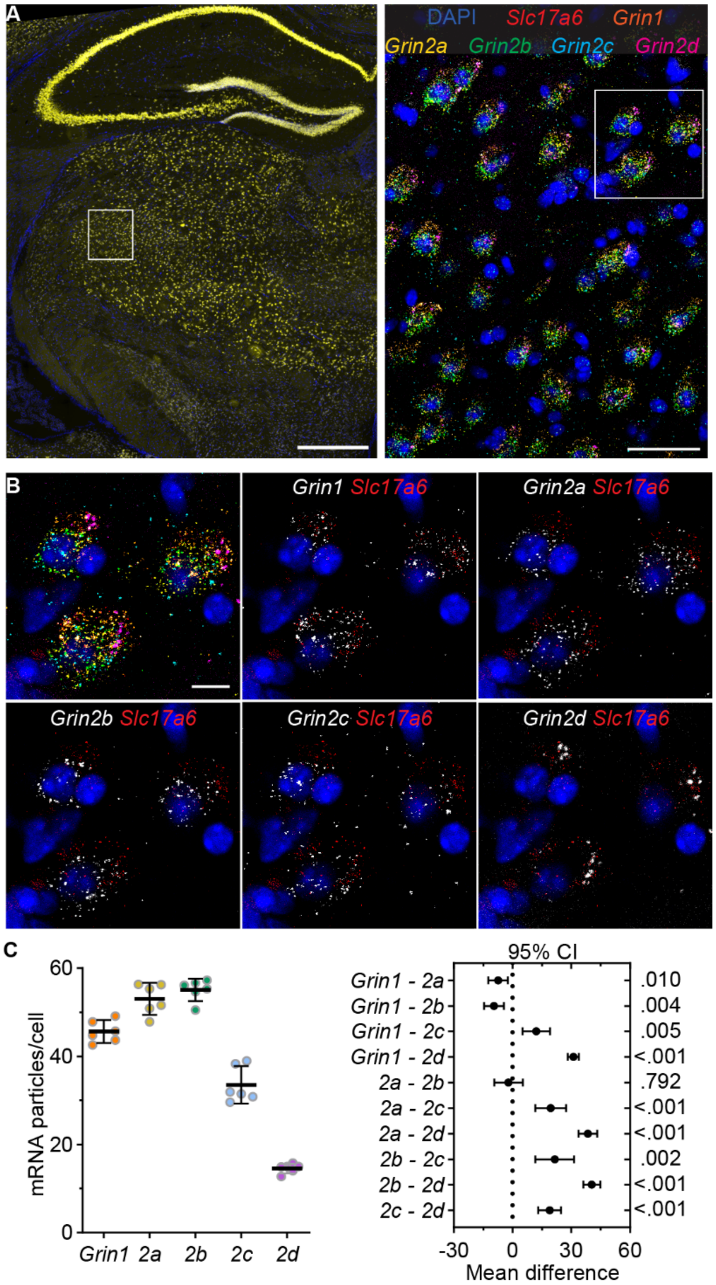


**Supplemental Figure S2. GluN subunit mRNA expression levels differ in the VPM thalamus. A.** Tiled 40x image of a fixed mouse brain section shows RNAscope detection of *Grin2a* mRNA (yellow) and DAPI-stained nuclei (blue). **B.** The boxed region from panel A shows a 40x image of *Grin1, Grin2a, Grin2b, Grin2c, Grin2d,* and *Slc17a6* (red) mRNA with DAPI. **C.** Images of the boxed region from panel B depict the merged signals as well as each *Grin* gene along with *Slc17a6* mRNA and DAPI. **D**. The plot shows the mean mRNA particle number per neuron (averaged across approx. 30 neurons per mouse) for 6 mice with the group mean ± *SD*. Data were analyzed by one-way RM-ANOVA, $F$ (2.059, 10.29) = 225.3, $p < 0.001$. At right, the plot shows the 95% CI of the mean difference for pairwise comparisons between all *Grin* genes with Tukey's test $p$ values listed along the right y-axis. Scale bars: (A) 500 µm, (B) 40 µm, and (C) 10 µm.

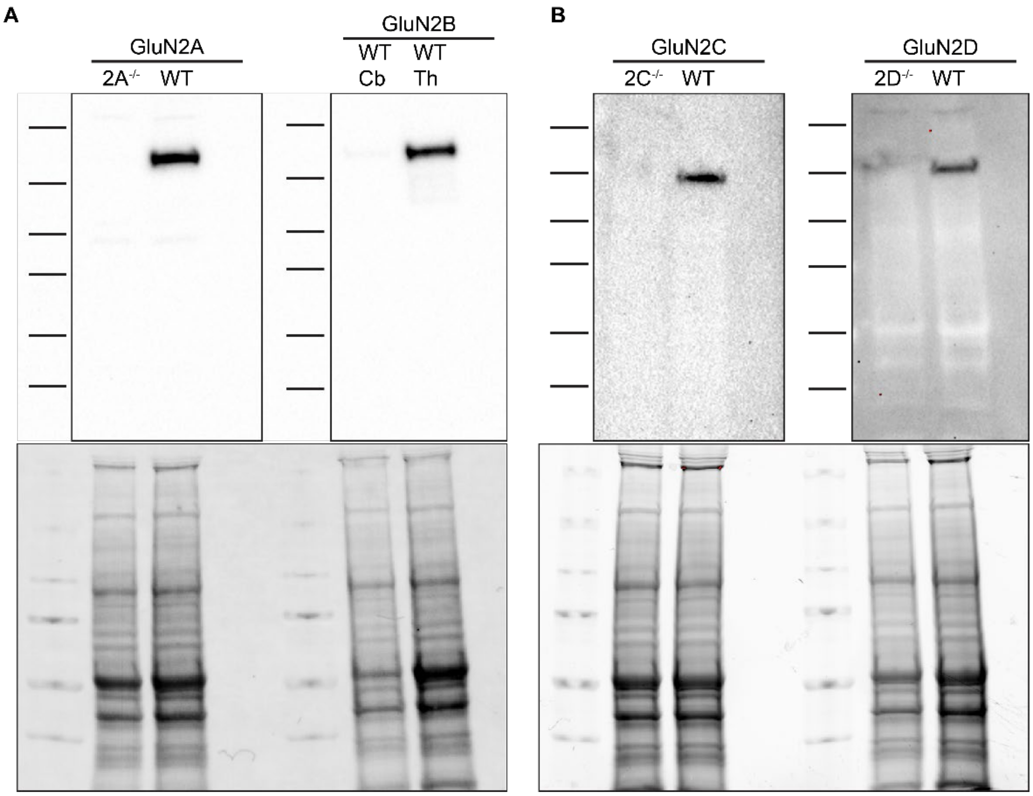


**Supplemental Figure S3. Validation of GluN2 antibodies for western blotting. A.** At left, whole thalamus from WT and *Grin2a* null ($2A^{-/-}$) mice were homogenized and separated by SDS PAGE, and then immunoblotted with GluN2A antibodies. At right, whole thalamus (Th) from 4-week-old WT mice and whole cerebellum (Cb) from 12-week-old mice were homogenized and separated by SDS PAGE, and then immunoblotted with GluN2B antibodies. **B**. Thalamus from WT mice as well as *Grin2c* null ($2C^{-/-}$) mice (left) and *Grin2d* null ($2D^{-/-}$) mice were processed as in panel A, and then immunoblotted with GluN2C and GluN2D antibodies, respectively. Total protein signals from each gel are shown at the bottom of panels A and B.

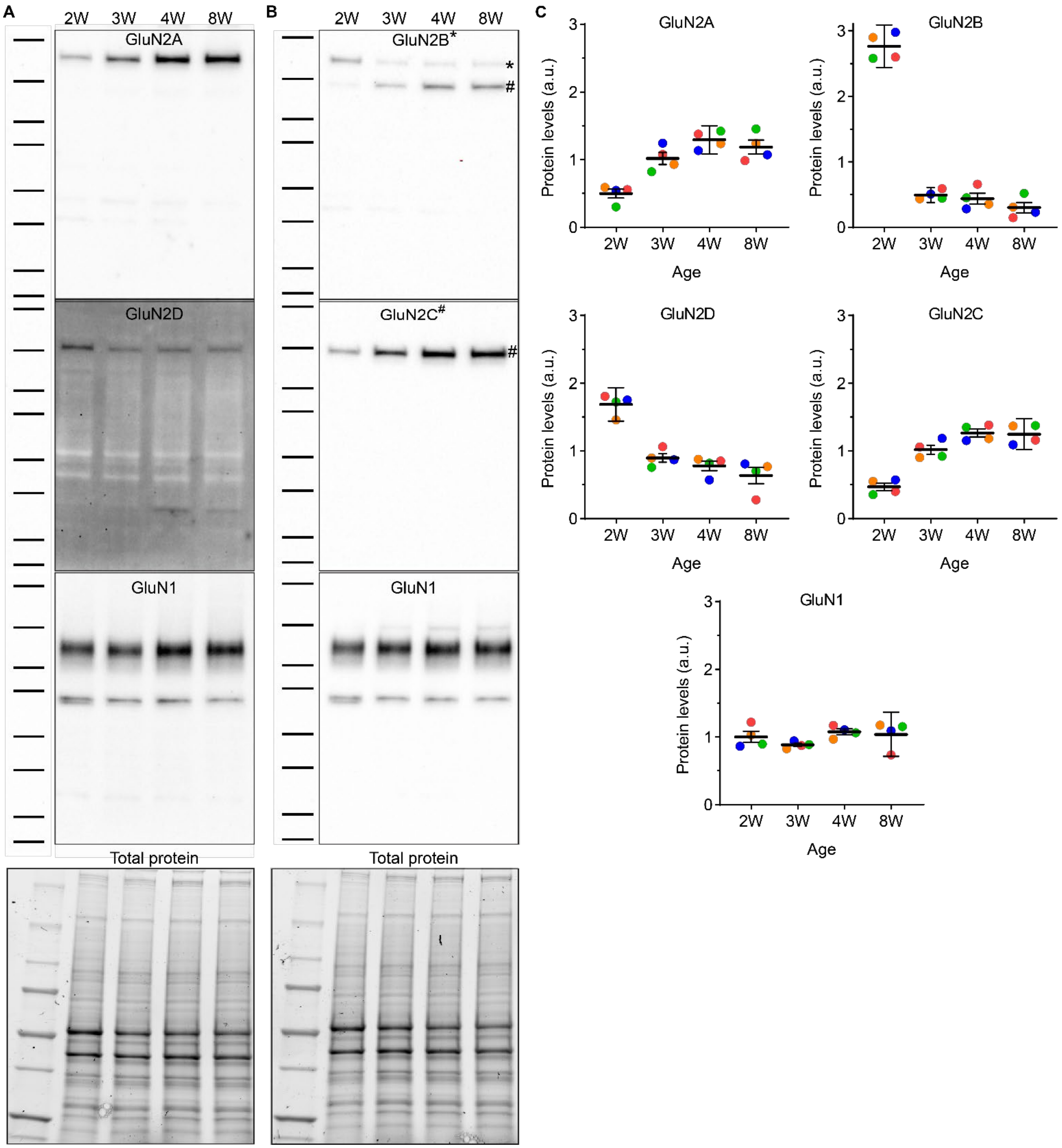


**Supplemental Figure S4. Developmental expression of NMDAR subunits in the cerebellum.** Whole cerebellum from mice aged 2, 3, 4, and 8 weeks (W) were homogenized, processed for SDS PAGE, run on duplicate gels, and then samples were immunoblotted. Samples were probed for (**A**) GluN2A, GluN2D, and GluN1, and (**B**) GluN2B, GluN2C, and GluN1. In the top image of panel B, the higher band (*) is GluN2B and the lower band is residual GluN2C (#) not removed by stripping buffer prior to probing for GluN2B. Protein levels were quantified by densitometry and normalized to GluN1 levels as well as total protein within each gel. **C**. Protein levels for each replicate ($n = 4$ mice; 2F, 2M) were shown as individual data points of the same color across age and expressed as fold change from the mean intensity across all ages to control for differences in band intensity across experiments. The line and error bars are mean ± SD.

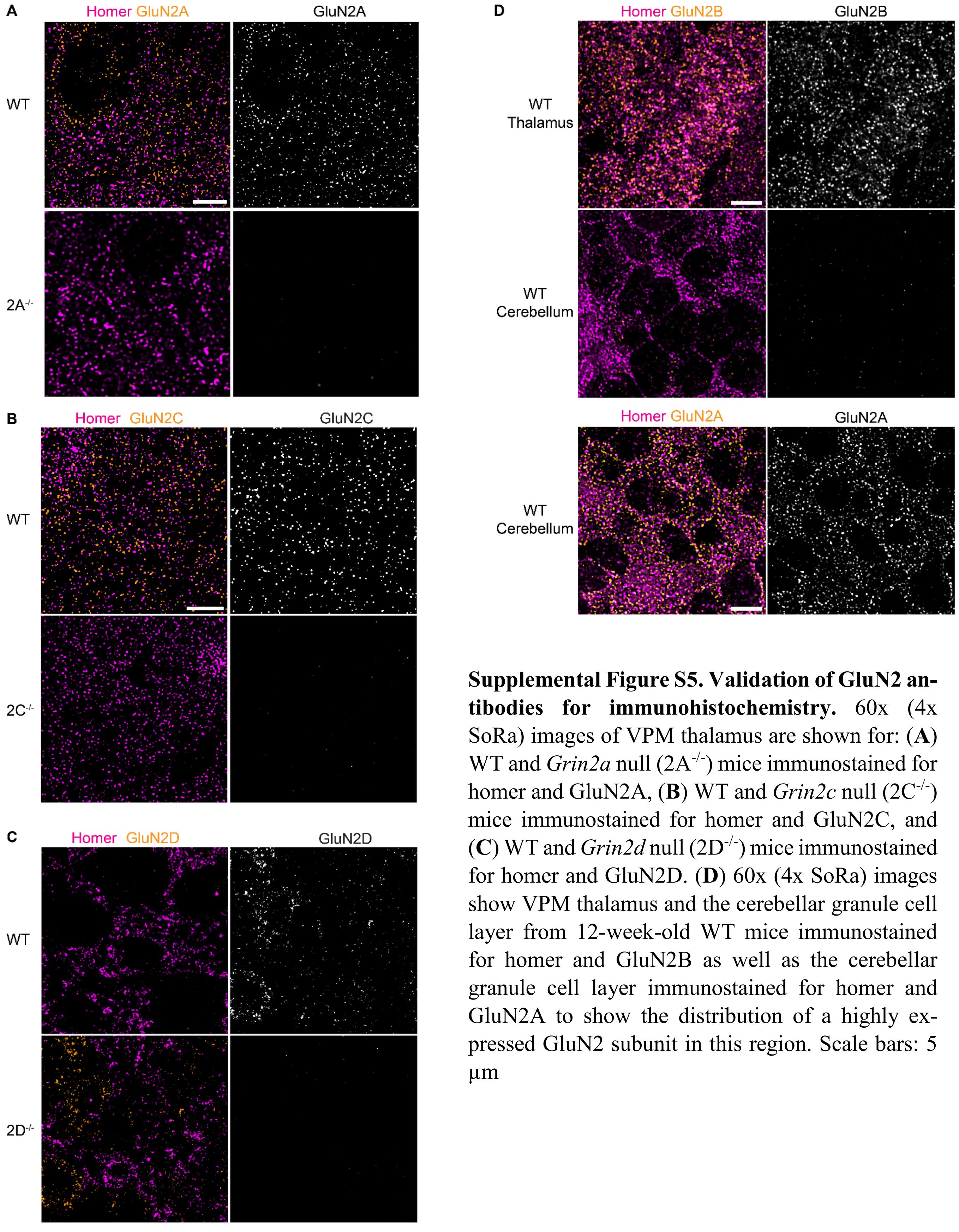


**Supplemental Figure S5. Validation of GluN2 antibodies for immunohistochemistry.** 60x (4x SoRa) images of VPM thalamus are shown for: (**A**) WT and *Grin2a* null ($2A^{-/-}$) mice immunostained for homer and GluN2A, (**B**) WT and *Grin2c* null ($2C^{-/-}$) mice immunostained for homer and GluN2C, and (**C**) WT and *Grin2d* null ($2D^{-/-}$) mice immunostained for homer and GluN2D. (**D**) 60x (4x SoRa) images show VPM thalamus and the cerebellar granule cell layer from 12-week-old WT mice immunostained for homer and GluN2B as well as the cerebellar granule cell layer immunostained for homer and GluN2A to show the distribution of a highly expressed GluN2 subunit in this region. Scale bars: 5 µm

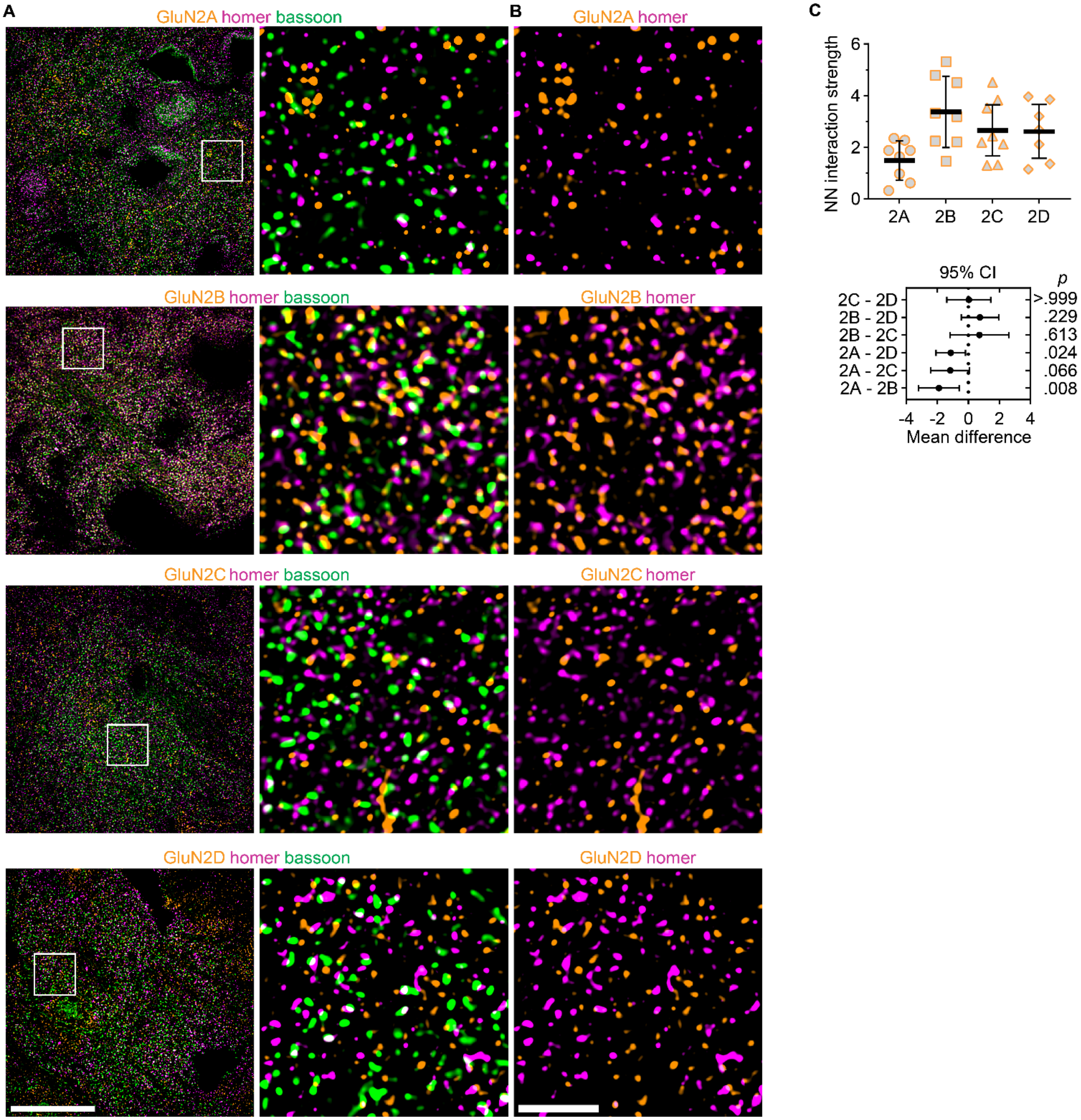


**Supplemental Figure S6. GluN2-homer nearest neighbor analysis. A**. 60x (4x SoRa) images show GluN2, homer, and bassoon immunolabeling with the boxed region shown at right. **B**. GluN2 and homer signals are shown to highlight their relative spatial distributions. **C.** NN interactions strengths were plotted for each mouse with group mean ± *SD*. Groups were compared by a mixed-effects model and Tukey's pairwise comparison tests for which the 95% CI of the mean difference and *p* values are shown beneath the group data. $F$ (1.89, 12.6) = 7.08, $p$ = 0.009, n = 8 mice (4F, 4M). Scale bars: A) 20 µm; B) 3 µm.

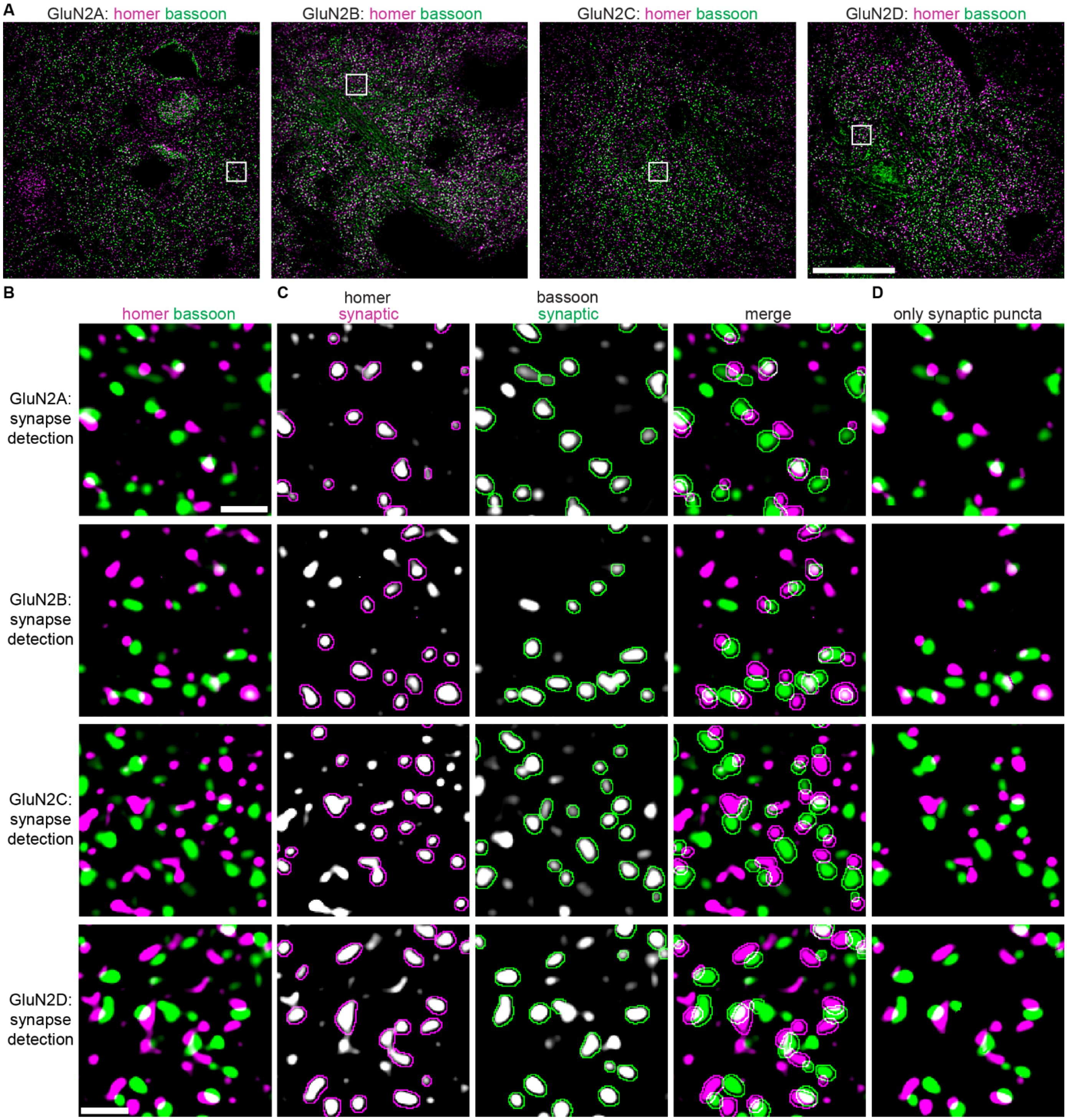


**Supplemental Figure S7. Homer-bassoon synapse detection for synaptic GluN2 localization (**Figure 3**). A.** 60x images (4x SoRa) show homer and bassoon puncta from respective GluN2A – GluN2D images in Figure 3. **B.** The boxed regions from panel A show individual homer and bassoon puncta. **C**. Homer and bassoon signals from panel B are shown separately in grayscale and merged with outlines depicting all detected postsynaptic homer and presynaptic puncta. The thresholded "only synaptic puncta" image shows only the outlined bassoon and homer puncta. Scale bars: A) 20 µm; B) 1 µm.

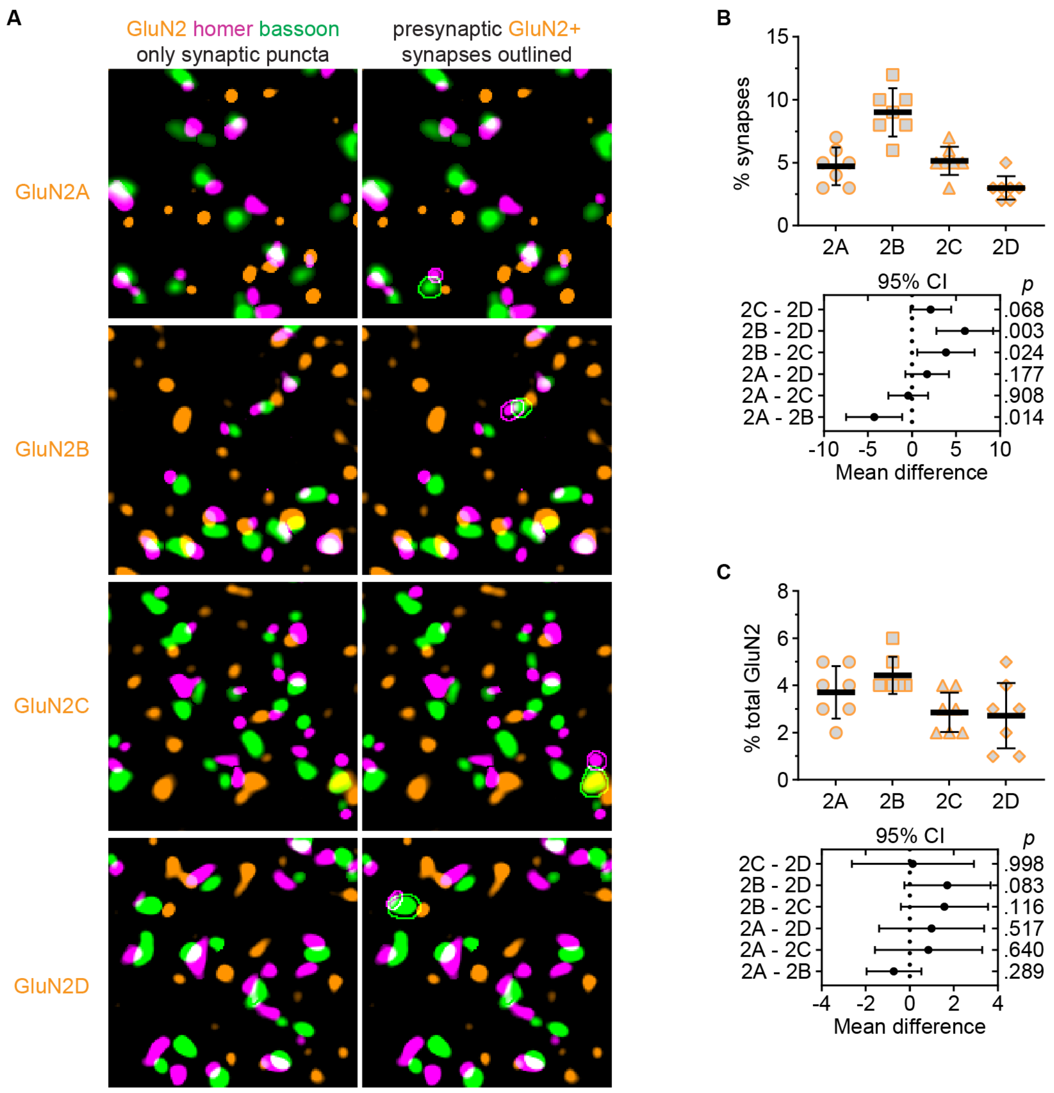


**Supplemental Figure S8. Presynaptic GluN2 localization. A.** 60x (4x SoRa) images show GluN2A – GluN2D immunolabeling with synaptic homer and bassoon puncta (same regions as shown in Figure 3). At right, synaptic puncta with GluN2 overlapping presynaptic bassoon without overlapping postsynaptic homer are outlined. **B**. The percentage presynaptic bassoon overlapping GluN2 and **C**) the percentage of GluN2 overlapping presynaptic bassoon were quantified in the full FOV, averaged across three images per mouse, and plotted with group mean ± *SD* (n = 7 mice; 4F, 3M). Data were analyzed with a mixed effects model and Tukey's pairwise comparisons for which the 95% CI fo the mean difference are plotted beneath the group data with the *p* values on the right y-axis. **B:** $F$ (2.52, 20.2) = 21.4, $p < 0.001$, n = 7 (4F, 3M); **C**) $F$ (2.25, 18.0) = 3.65, $p = 0.042$.

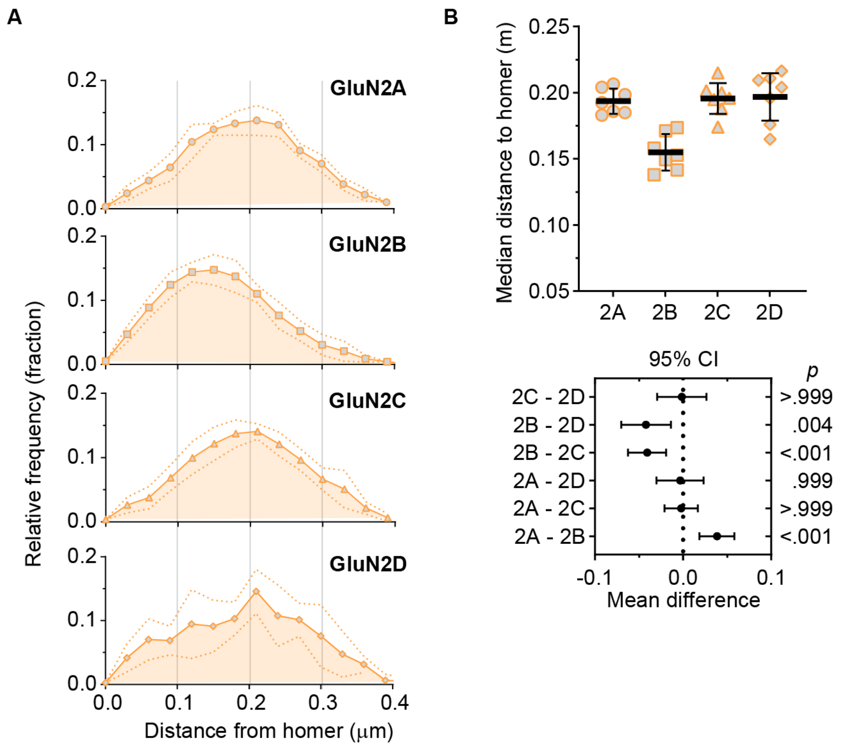


**Supplemental Figure S9. GluN2B is localized closest to postsynaptic homer. A**. The distances between the maxima of overlapping postsynaptic homer and GluN2 puncta were measured and plotted as a frequency distribution for each mouse (n = 7 mice; 4F, 3M). The plot shows the group mean frequency distribution ± *SD*. **B.** The median distance was determined for each mouse and plotted with group mean ± *SD*. The data were analyzed by Brown-Forsythe ANOVA, $F$ (3, 18.9) = 14.2, $p < 0.001$, with pairwise Dunnett T3 comparisons for which the 95% CI of the mean difference is plotted with $p$ values on the right y-axis.

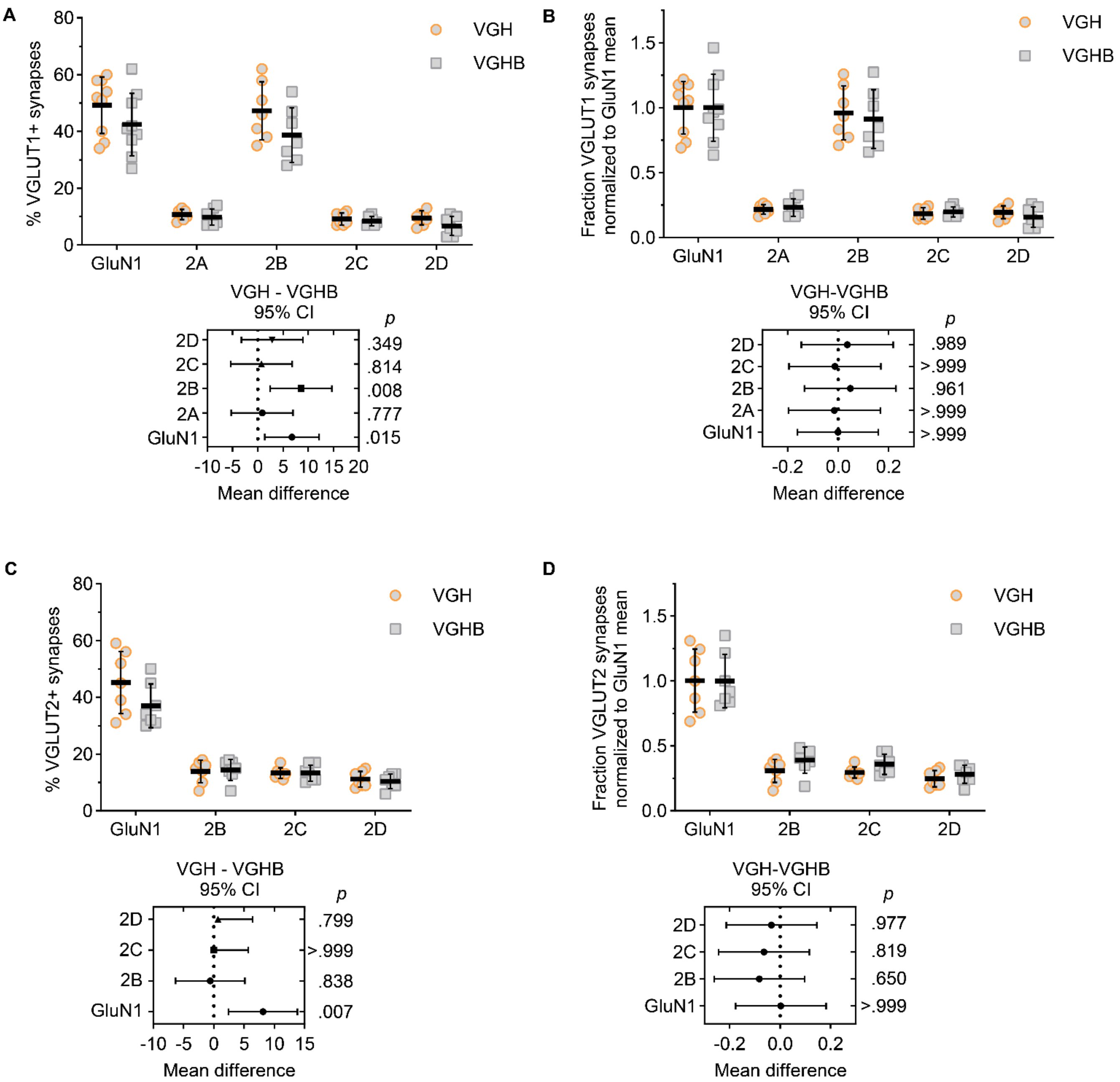


**Supplemental Figure S10. Comparison of fraction VGLUT-positive synapses with GluN with and without bassoon. A.** The fraction of VGLUT1-positive synapses with GluN2 overlapping homer was measured for immunostainings of VGLUT1, GluN, and homer (VGH) or VGLUT1, GluN, homer, and bassoon (VGHB). VGH defined postsynaptic homer as those homer puncta overlapping VGLUT1, while VGHB defined postsynaptic homer as homer puncta within homer-bassoon pairs with bassoon overlapping VGLUT1. The percentages of synapses with GluN1 and GluN2B significantly differed between these two methods as determined by a mixed-effects model (main effects, GluN: F (1,32) = 9.07, p = .005; interaction: F (4, 32) = 1.46, p = .237) with posthoc Sidak's tests comparing VGH and VGHB values for each GluN subunit. **B**. The values in panel A were normalized to the GluN1 mean within each group (VGH and VGHB) and the data were analyzed by mixed-effects model to determine if the relative differences between GluN subunits differed between the two methods (main effect, GluN: F (1, 32) = 0.147, *p* = 0.704; interaction: F (4, 32) = 0.192, *p* = 0.941; n = 7 mice; 4F, 3M). There were no significant differences between the relative values for each GluN subunit. VGLUT2 data shown in panels C and D were analyzed as described for VGLUT1 data. GluN2A was not included because four-channel imaging was not possible with our antibody combinations. **C**. Mixed-effects model, main effect, GluN: F (1, 24) = 2.24, *p* = .148; interaction: F (3, 24) = 2.17, *p* = .117. **D.** Mixed-effects model, main effect, GluN: F (1, 24) =

1.76, $p$ = .197; interaction: F (3, 24) = 0.312, $p$ = .817; n = 7 mice; 4F, 3M. There were no significant differences in the relative values for each GluN subunit.

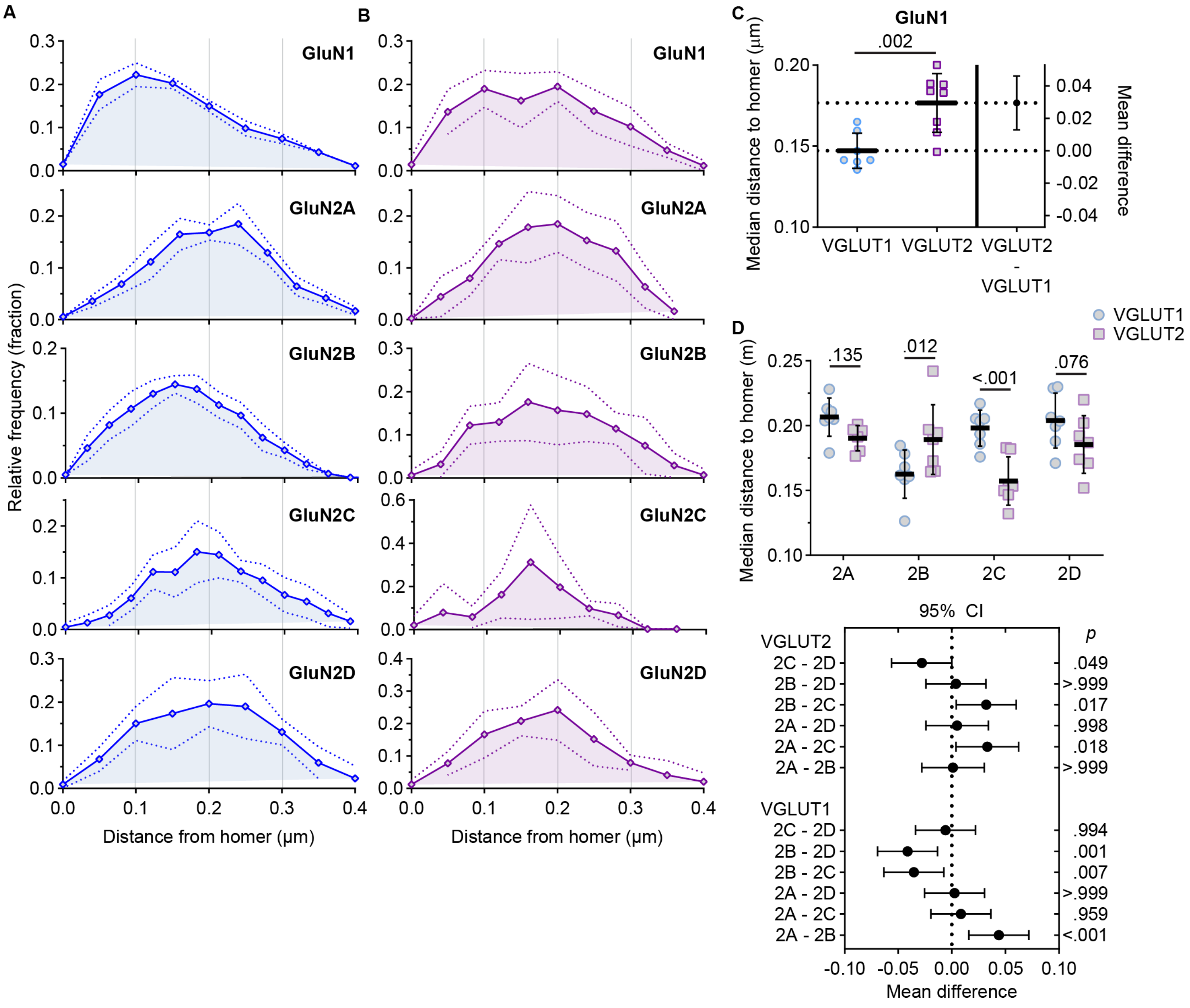


**Supplemental Figure S11. NMDAR distance from the synapse is input- and GluN2 subtype-specific.** The distances between the maxima of overlapping postsynaptic homer and GluN2 puncta were measured for **A**) VGLUT1- and **B**) VGLUT2-positive synapses and plotted as a frequency distribution for each mouse. The plots show the group mean frequency distribution ± *SD* for each GluN subunit. **C.** GluN1 median distance for each mouse was plotted with group mean ± *SD* (VGLUT1: n = 7 mice; 4F, 3M; VGLUT2 = n = 8; 4F, 4M) on the left y-axis of an estimation plot with the mean difference and 95% CI plotted on the right y-axis. Groups were compared by a Welch's t-test ($t$ = 3.88, df = 11.7). **D**. The median distance was determined for each GluN2 subunit and plotted with group mean ± *SD*. The data were analyzed by two-way ANOVA with pairwise Sidak comparisons for which the 95% CI of the mean difference is plotted with $p$ values on the right y-axis. Interaction: $F$ (3, 47) = 7.64, $p$ < 0.001; GluN2 effect: $F$ (3, 47) = 4.97, $p$ = 0.004; VGLUT effect: $F$ (1, 47) = 5.62, $p$ = 0.022.

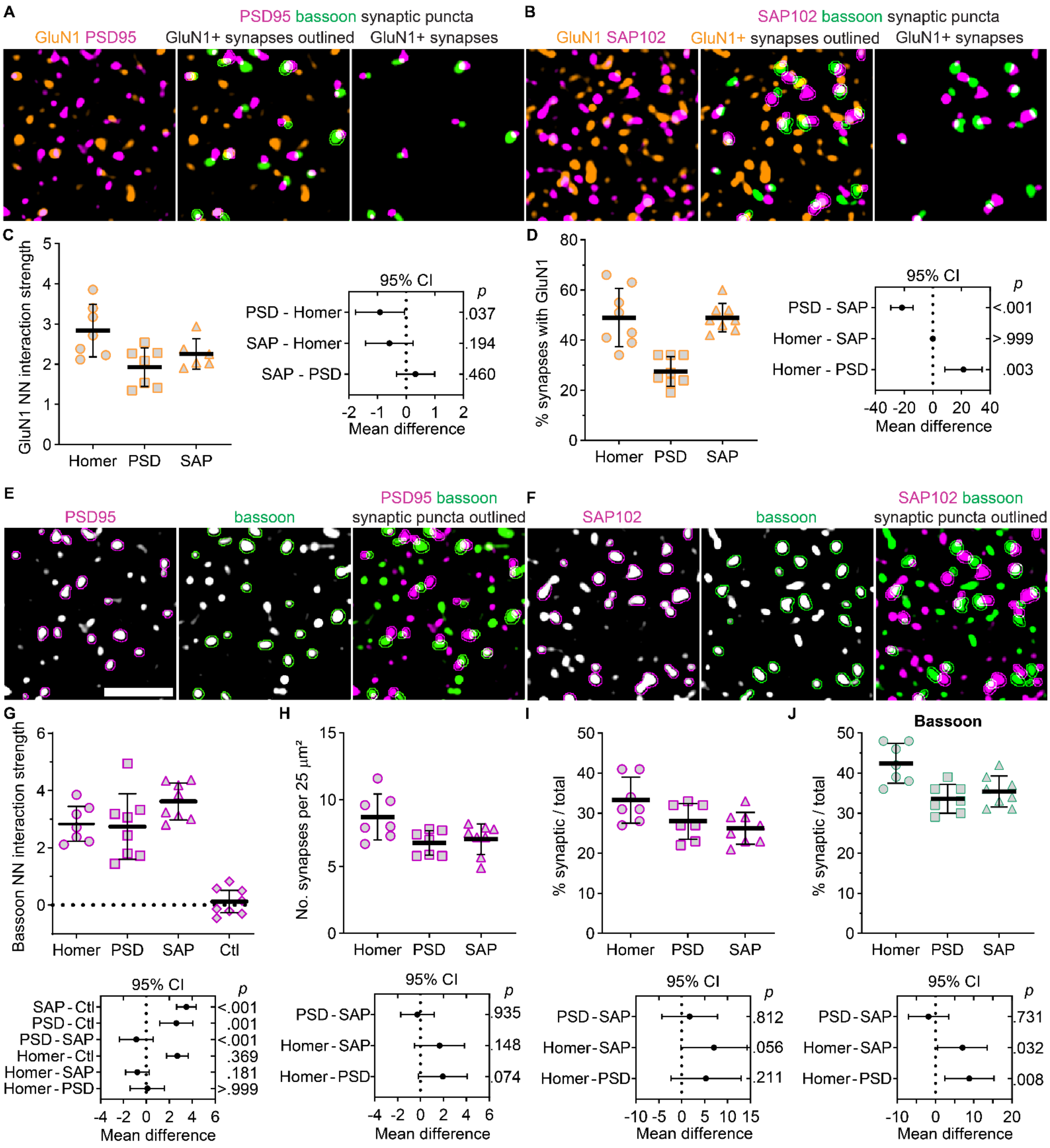


**Supplemental Figure S12. Comparison of synapse detection and GluN1 colocalization with homer, PSD95, and SAP102.** 60x (4x SoRa) images show GluN1 with postsynaptic markers (**A**) PSD95 or (**B**) SAP102, GluN1 with synaptic puncta and GluN1-positive synapses outlined, and only GluN1-positive synapses. **C**. The NN interaction strength for GluN1 with homer, PSD95, and SAP102 and (**D**) the percentage of synapses with GluN1 overlapping the postsynaptic marker were plotted with the group ± *SD*. The data were analyzed by mixed-effects models with pairwise Tukey's comparisons for which the 95% CI of the mean difference is plotted (at right) with *p* values on the right y-axis. C: F (2, 14.95) = 5.593, *p*

= 0.015. D: $F (2, 13.8) = 18.3$, $p < 0.001$. 60x (4x SoRa) images show (**E**) PSD95 and (**F**) SAP102 immunostaining with bassoon in grayscale and the merge with the puncta detected as synapses outlined. The following data were plotted with the group mean ± *SD*, and analyzed by mixed effects models and Tukey's pairwise comparisons for which the 95% CI of the mean difference is plotted (bottom) with *p* values on the right y-axis: **G**) NN interaction strength for bassoon with homer, PSD95, and SAP102 as well as bassoon with homer signal rotated 90 degrees ($F (3, 18.4) = 31.2$, $p < 0.001$), **H**) the number of synapses detected as pairs of bassoon with homer, PSD95, and SAP102 per 25 $\mu m^2$ ($F (2, 14.1) = 4.56$, $p = 0.030$), **I**) the percentage of total homer, PSD95, and SAP102 puncta detected as synaptic ($F (2, 16.5) = 4.24$, $p = 0.033$), and **J**) the percentage of total bassoon puncta detected as synaptic when paired with homer, PSD95, and SAP102 ($F (2, 17.0) = 8.79$, $p = 0.002$).
